# Auto-qPCR: A Python-based web app for automated and reproducible analysis of qPCR data

**DOI:** 10.1101/2021.01.14.426748

**Authors:** Gilles Maussion, Rhalena A. Thomas, Iveta Demirova, Gracia Gu, Eddie Cai, Carol X.-Q Chen, Narges Abdian, Theodore J.P. Strauss, Sabah Kelaï, Angela Nauleau-Javaudin, Lenore K. Beitel, Nicolas Ramoz, Philip Gorwood, Thomas M. Durcan

**Author notes:** Corresponding author: Thomas M. Durcan, Ph.D. McGill University, The Neuro’s Early Drug Discovery Unit (EDDU) 3801 University St., Montreal, QC H3A 2B4 Canada Mail. These authors have contributed equally (Gilles Maussion, Rhalena A. Thomas). Abbreviations: CT (cycle threshold), qPCR (quantitative polymerase chain reaction), iPSC (induced pluripotent stem cells), CNVs (copy number variants), SNVs (single nucleotide variants), DA (dopaminergic), Neural Precursor Cells (NPC), DA neurons (DANs).

## Abstract

Quantifying changes in DNA and RNA levels is essential in numerous molecular biology protocols. Quantitative real time PCR (qPCR) techniques have evolved to become commonplace, however, data analysis includes many time-consuming and cumbersome steps, which can lead to mistakes and misinterpretation of data. To address these bottlenecks, we have developed an open-source Python software to automate processing of result spreadsheets from qPCR machines, employing calculations usually performed manually. Auto-qPCR is a tool that saves time when computing qPCR data, helping to ensure reproducibility of qPCR experiment analyses. Our web-based app (https://auto-q-pcr.com/) is easy to use and does not require programming knowledge or software installation. Using Auto-qPCR, we provide examples of data treatment, display and statistical analyses for four different data processing modes within one program: (1) DNA quantification to identify genomic deletion or duplication events; (2) assessment of gene expression levels using an absolute model, and relative quantification (3) with or (4) without a reference sample. Our open access Auto-qPCR software saves the time of manual data analysis and provides a more systematic workflow, minimizing the risk of errors. Our program constitutes a new tool that can be incorporated into bioinformatic and molecular biology pipelines in clinical and research labs.

## Introduction

Polymerase chain reaction (PCR) identifies a nucleic acid fragment of interest by increasing its proportion relative to others ^1^. Initially the technique was primarily used to visualize DNA fragments for cloning ^2, 3^ or genotyping ^4–6^, but can now be used to investigate genetic polymorphisms and mutations ^7, 8^, copy number variants (CNVs) ^9^, single nucleotide variants (SNVs), point mutations, and genetic deletion/duplication events ^10^. With the development of fluorogenic probes and dyes capable of binding newly synthesized DNA, PCR became more quantitative, leading to innovative tools for quantifying relative transcript levels for one or more genes, now referred to as quantitative PCR (qPCR). With these technological advancements, qPCR is now used to quantify messenger RNA (mRNA) ^11^, long non-coding RNA ^12^, microRNAs ^13,14^DNA-protein interactions ^15^ and epigenetic modifications ^16, 17^. Thus, the advent of PCR has revolutionized our ability to analyze and quantify nucleic acids and has made qPCR a standard technique.

qPCR experiments are already automated at the data acquisition stage, with thermocycler software providing “by default” pre-processing procedures ^18^. However, several steps (data exclusion, normalization, data display and differential analyses) required for full data interpretation are heterogenous, and the data processing and display methods and options vary widely across available licenced qPCR programs. Commercially available software that provide data summaries and statistical output do not systematically allow for user selections and are not necessarily transparent as to the processes and settings being used. Also, not all qPCR software provides a statistical output. Analysis of qPCR data is still highly time consuming and error prone, especially when processing large numbers of data points. The user must intervene to include or exclude replicates, which, without guidelines or standardized procedures, can potentially introduce “user-dependent” variation and errors. To both simplify and accelerate this data analysis step for qPCR datasets, we have created a Python-based, open source, user-friendly web application “Auto-qPCR” to process exported qPCR data and to provide summary tables, visual representations of the data, and statistical analysis. The program can be found at the website https://auto-q-pcr.com/.

The program can work with the two commonly used molecular biology approaches: (i) absolute quantification, where all RNA estimations rely on orthogonal projection of the samples of interest onto a calibration curve ^19^, and (ii) relative quantification that relies on difference of cycle threshold (CT) values between the gene of interest and endogenous controls ^20^.

Here we use Auto-qPCR to analyze qPCR datasets and illustrate four distinct computational methods.Overall, Auto-qPCR provides an all-in-one solution for the user, going from datasets to graphs, within one web-based software package. Unlike other software, the intermediate and final results are output by the program, allowing a full review of the data and accurate statistical treatment based on the experimental design. Auto-qPCR was conceived to build logical links between the experimental design and required statistics for differential analyses of each mode, which is rarely found in other qPCR programs. While other open-source qPCR analysis software programs and web apps ^21–23^ are available, they are only able to normalize, compare and display qPCR data generated with one of the two quantification modes ^19, 20^. In contrast, Auto-qPCR provides a comprehensive data analysis package for a wide variety of qPCR experiments. Using the web app does not require prior programming knowledge, account creation or desktop installation. Additionally, the program has been designed to assist the user at each step of the analysis once the exported data files have been collected from the qPCR system.

Auto-qPCR can be used to analyse qPCR data in a reproducible manner, simplifying data analysis, avoiding potential human error, and saving time. In this manuscript, we describe some of the uses of the software and outline the steps required, from entering an individual dataset to complete statistical analysis and graphical presentation of the data.

## Methods

### Culture of iPSC lines

To illustrate the four different models of quantification managed by the Auto-qPCR program, we used 11 different iPSC cells lines whose properties are presented in (**Table S1)**. Quality control profiling for the iPSCs used was outlined previously ^24^.

The iPSCs were seeded on Matrigel-coated dishes and expanded in mTESR1 (Stemcell Technologies) or Essential 8 (ThermoFisher Scientific) media.Cells were seeded at 10 to 15% confluency and incubated at 37°C in a 5% CO2 environment. The media was changed daily until the cultures reached 70% confluency. Cells harbouring irregular borders, or transparent centres were manually removed from the dish prior to dissociation with Gentle Cell Dissociation media (Stemcell Technologies). The IPSCs were then seeded and differentiated into cortical or dopaminergic neuronal progenitors or neurons.

### Generation of cortical and dopaminergic neurons

The induction of cortical progenitors was performed as described previously ^25^. The media used for cortical differentiation is described in the standard operating procedure published on the Early Drug Discovery Unit (EDDU) website ^24^. Once neural progenitor cells (NPCs) attained 100% confluency, they were passaged and seeded on a Poly-Ornithine-laminin coated dishes to be differentiated into neurons. Cells were switched for 24 hours to 50% Neurobasal (NB) medium, and 24 hours later placed in 100% NB medium with AraC (0.1µM) (Sigma) to reduce levels of dividing cells. After the third day of differentiation, cells were maintained in 100% NB medium without AraC for four days before being collected for RNA extraction. IPSCs were induced into dopaminergic NPCs (DA-NPCs) according to methods previously described ^26^, modified according to methods used within the group ^27^. DA-NPCs were subsequently differentiated into dopaminergic neurons (DANs), with immunostaining and qPCR analysis performed at four and six weeks of maturation from the NPC stage ^28^.

### DNA and RNA extraction

IPSCs were dissociated with Gentle Cell Dissociation Reagent (Stem Cell Technologies) while Accutase® Cell Dissociation Reagent (Thermo Fisher Scientific) was used to dissociate NPCs and iPSC-derived neurons. After 5 minutes incubation at 37°C with the indicated dissociation agent, cells were collected and harvested by centrifugation for 3 minutes at 1200 rpm. Cell pellets were resuspended in lysis buffer and stored at -80°C before DNA or total RNA extraction with the Genomic DNA Mini (Blood/Culture Cell) (Genesis) or mRNAeasy (Qiagen) kits, respectively.

### cDNA synthesis, quantitative PCR, and data export

Reverse transcription reactions were performed on 400ng of total RNA extract to obtain cDNA in a 40μl total volume containing, 0.5μg random primers, 0.5mM dNTPs, 0.01M DTT and 400 U/µl-MMLV RT (Carlsbad, CA, USA). The reactions were conducted in singleplex, in a 10µl total volume containing 2X Taqman Fast Advanced Master Mix, 20X Taqman primers/probe set (Thermo Fisher Scientific), 1µl of diluted cDNA and RNAse-free H2O. Real-time PCR (RT-PCR) was performed on a QuantStudio 3 machine (Thermo Fisher Scientific). Primers/probe sets from Applied Biosystems were selected from the Thermo Fisher Scientific web site . Two endogenous controls (beta-actin and GAPDH) were used for normalization (**Table S2)**.

Data generated from the QuantStudio machine were extracted using QuantStudio design and analysis software, either (i) as excel files (*.xls or *.xlsx extensions) and the results tab was saved as a ‘comma delimited’ csv file or (ii) extracted as a txt file that only contained the result tab..

### Collection of external data set

An external qPCR data set was provided from an earlier published study ^29^, which quantified levels of *Nrxns* and *Nlgn* transcripts in the subcortical areas of the brains from mice submitted to conditioned place preference (CPP) with cocaine. Briefly, subcortical areas (subthalamic nucleus, globus pallidum and substantia nigra) of sectioned mouse brains were isolated by laser capture microdissection. RNA was extracted with the Arcturus PicoPure kit and reverse transcription performed as above. The qPCR experiments were performed according to an absolute quantification design on the Opticon 2 PCR machine (Biorad). Β2Microglobulin (*B2M*) was used as endogenous control. Data were re-extracted from the Opticon Monitor 2 files as csv files and analyzed by Auto-qPCR.

### Program development and structure

The program was written in Python using Pandas and NumPy. A main script calls the selected model script (absolute.py, relative.py and stability.py), which processes the data and then calls the statistical functions script (if selected) and the plotting function script. The graphical user interface (GUI) was created using Flask, a package for integrating HTML and Python code. The GUI is written in JavaScript, CSS, HTML and Bootstrap4, a framework for building responsive websites. Our GitHub repository (https://github.com/neuroeddu/Auto-qPCR) includes all python processing scripts and scripts to build the GUI that can be installed locally to run on a computer. A complete list of package dependencies and instructions to install and run the package app locally are posted in the GitHub repository. The program was developed using git version control with multiple contributors. The web app is hosted by the Brain Imaging Centre at the Montreal Neurological Institute-Hospital (The Neuro) and was installed in a virtual machine directly from the public GitHub repository. When updates are available the changes will be applied to the web app using GitHub. The organization and function of the script files for the program are in **Tabel 3**. The web app can be found at https://auto-q-pcr.com (**Figure S1**)

### Program function - input data processing and quantification

The Auto-qPCR program reads the raw data in the form of a results spreadsheet (via the users file navigator) and reformats it into a data frame in Python. The user enters information into the web app read as arguments by the software. See **Table S4** for a list of all the user inputs and **Figure S2** for examples of the input files. The values for the reference genes/targets (*ACTB*, *GAPDH*) are calculated for each sample and technical replicate (cell line, time point, treatment condition) separately.

To detect outliers, the standard deviation (std) of the technical replicates for a given sample is calculated, if the std is greater than the cut-off (the default value is 0.3), then the technical replicate furthest from the sample mean is removed. The process occurs recursively until the std is less than the cut-off or the value of “max outliers” is reached. The 0.5 default means that outliers will be removed until two technical replicates remain. The ‘preserve highly variable replicates’: If the CT-std is less than 0.3, but the absolute (mean-median)/median is less than 0.1, replicates are preserved. This helps to account for a lack of a clear outlier, where two of three replicates are close to equally distributed around the mean.

Model dependent processing: **Absolute model** calculates the ratio between the gene of interest and each control. For each gene/target of interest the normalized value is calculated against the mean of each control target separately, then the mean value from normalized to controls is calculated. **Relative model ΔCT**, without a calibration sample, calculates the ΔCT by subtracting the Control CT value from the CT value for the target from each (endogenous control), then takes mean value of the resulting deltas. **Relative model ΔΔCT** and **genomic stability model**, individually calculates the ΔCT for the target in test sample and the reference/calibration sample(s) then calculates the ΔΔCT by subtracting the reference ΔCT from the test sample. For all models, the mean value of technical replicates is calculated for each target.

For the relative models, values of reference genes are calculated separately for each input file. The data from one input file will not be applied to another file. For the absolute model, qPCR output for each gene is found in a separate file and the selected endogenous controls will be applied to all the data input in one analysis. For all models, two spreadsheets are created that can be opened in Excel.1) “clean_data.csv” contains the ΔCT calculated for each technical replicate, including outliers, indicated by “TRUE” in the column “Outlier”. 2) and “summary_data.csv” contains the mean, standard deviation (std) and standard error (SE) for each sample calculated from the included technical replicates; this output can easily be analyzed analyzed in another statistical program (R, SASS, Prism). All the input and output data are cleared after processing and no user data is stored in the web app.

### Program function – statistical analysis

For testing differential gene expression, the user selects the statistic option and files in a form to indicate the conditions of the experiment. Either paired test (t-test) or multiple comparisons (one-way ANOVA or 2-way ANOVA) to investigate interaction effects is selected. The names of the variables to be grouped by must be within either the ‘sample names’ column in the input file or within an additional column, which was created during the qPCR setup). A column can also be added manually into the results input file(s)file, although this will add a risk of copy/paste errors and add additional time to the analysis process. See **Table S5** for the list of which analysis is applied for each setting. All default setting are maintained for statistical functions (for details see the Pingouin documentation at https://pingouin-stats.org/, the output has been reformatted to be more easily read and interpreted by users and for consistency across statistical outputs.

### Program function – visualization

The plotting scripts were written using the Matplotlib bar chart function. The labels and axis settings were all adjusted directly within the script (plot.py). The user can dictate the gene/target order and the sample order (cell lines, treatments, time points) in the web app by entering the orders into the appropriate input box. The order variables will be grouped for the summary plots. All the plots are automatically generated and saved as png files. If statistics are applied, two summary bar charts of the mean values are generated, grouped by the selected variable. For two-way ANOVA analysis, the summary bar chart will group the first variable on the x-axis and the second variable will be visualized in different colours and indicated in the legend.

### Data availability and reproducibility

All raw csv input files data files and output files used in plots are available at https://github.com/neuroeddu/Auto-qPCR, along with a user guide. The example input (Input Data) and output files (Output Data) are all available and organized by Figure names. The parameters used for each figure can be found in the document “Notes_on_Datasets.docx” and screen shots of the filled web app from for each figure are in the Supplementary Figures. The example output will be replicated identically if the same conditions are entered.

### Illustrations

The schematic representation in **Figure 1** and simplified versions in **Figures 2-4** were created in Adobe Illustrator Creative Cloud 2020, with icons inserted from BioRender.

**Figure 1.**
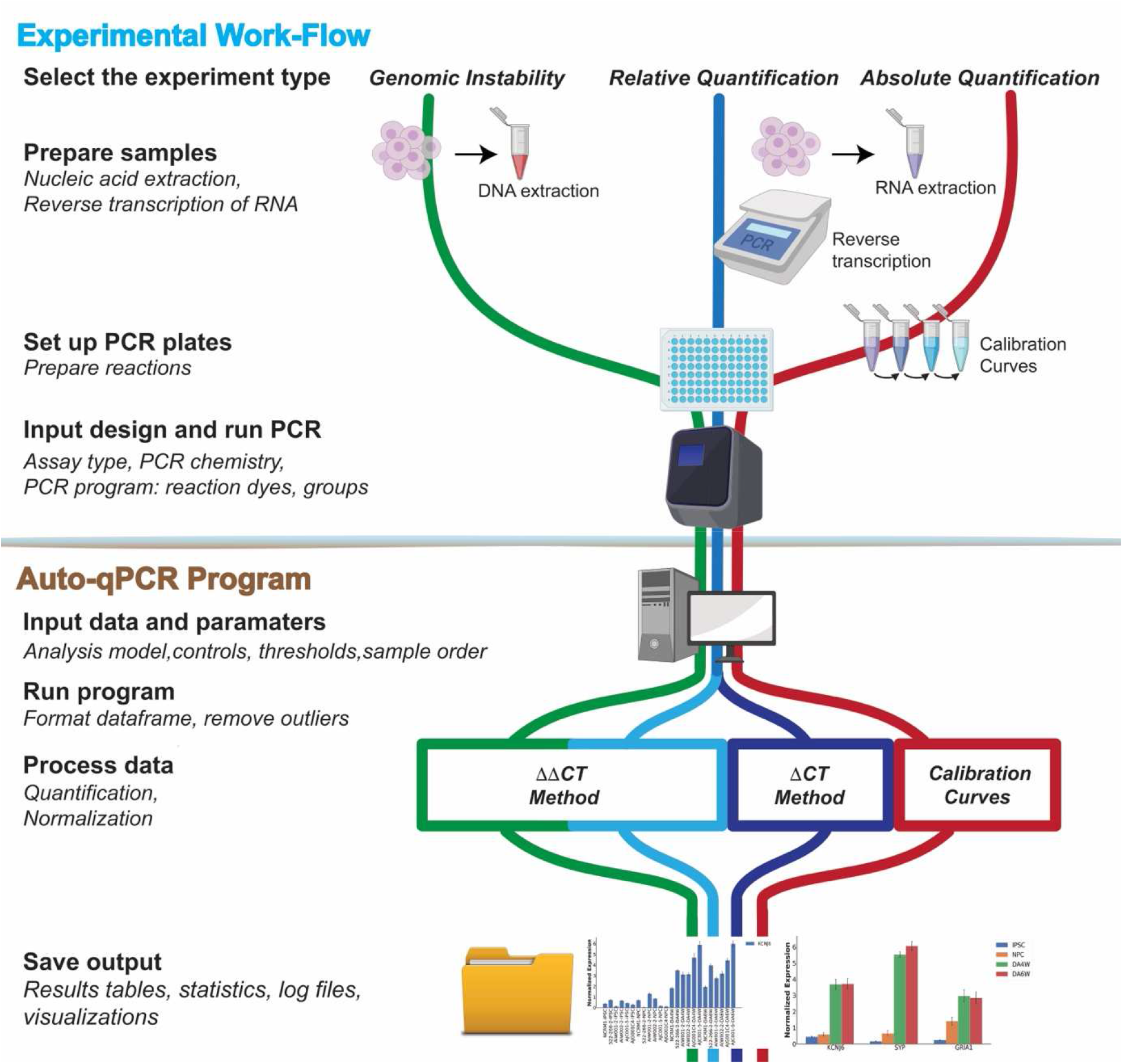
Workflow of a qPCR experiment. Schematic representation of common qPCR assays: genomic stability assay to detect DNA deletions or duplication events (green line), two methods to quantify RNA (cDNA) using either absolute (red line) or relative quantification designs (blue lines). qPCR experiments can be sub divided in two parts: the sample preparation and running the PCR machine (Experimental Workflow) and the data analyses (Auto-qPCR Program). The preparation of the experiment includes nucleic acid extraction followed by a cDNA synthesis step (for RNA) and the *in silico* design of the PCR plate layout. Nucleic acid preparations are accurately diluted. For the absolute model, a standard curve must be created. The experimental design of the PCR plate, including the chemistry (fluorophore, primer mix), the status of the samples, and the transcripts or DNA region that are going to be amplified, must be generated *in silico.* After having defined the parameters of the qPCR reactions (number of PCR cycles and length of the different steps (denaturation, hybridization and elongation), and the temperatures), the PCR is run. The exported data from the thermocycler, converted to csv, is entered into the Auto-qPCR software and the model matching the experimental design and parameters for analysis are selected. The software will reformat the data, quantify each sample normalized to controls, and create spreadsheets and graphs to visualize the data analyses, all of which will be included in a zip file for the user to save.

**Figure 2.**
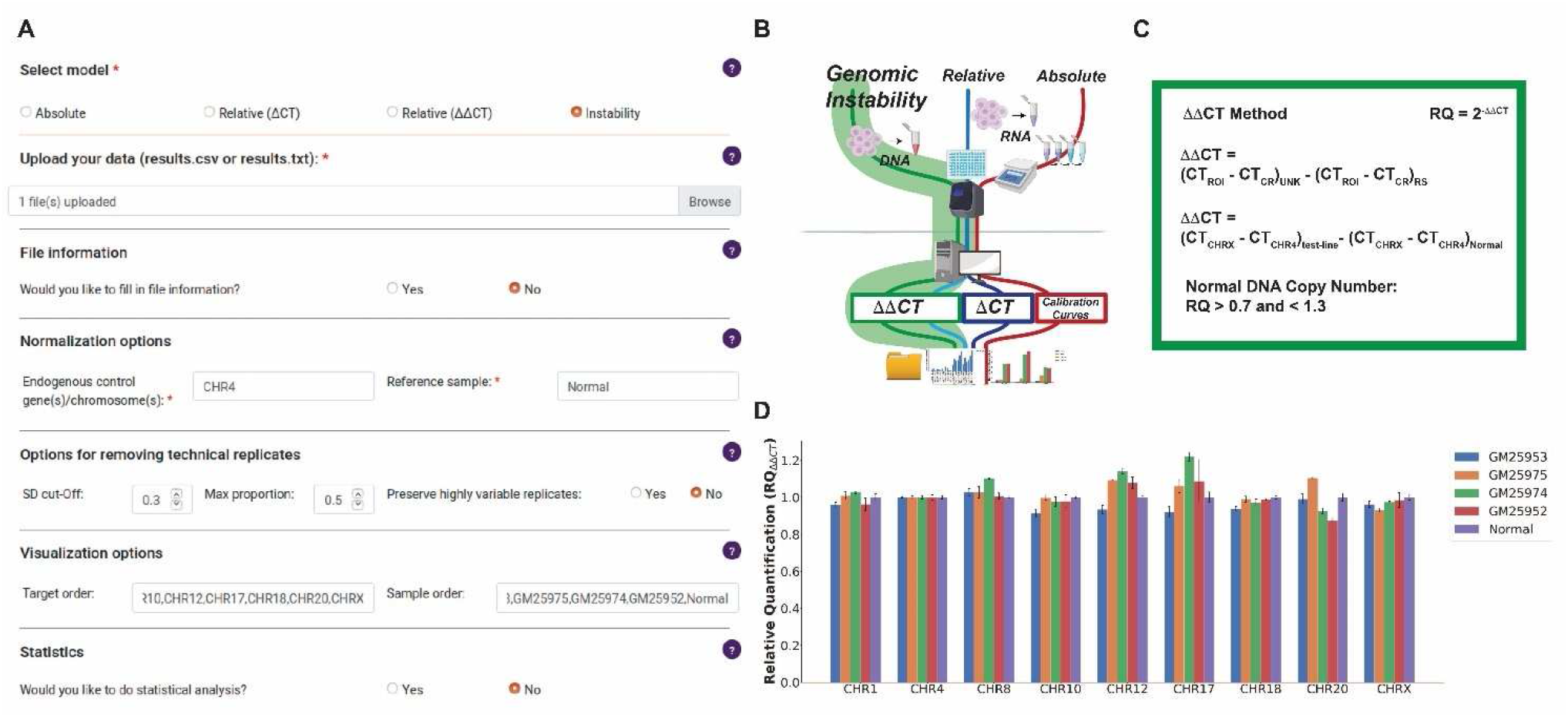
Auto-qPCR can process PCR genomic stability data. (**A**) Screen capture of the Auto-qPCR web-app. (**B**) Simplified schematic of PCR workflow showing the genomic instability analysis in green. The DNA copy number is quantified with the same formula as the ΔΔ CT relative quantification model. (**C**) The calculations carried out for genomic instability testing (ΔΔ CT). Top, the general formula used where the CT values for each chromosome were normalized to a region of interest and then to a reference sample. Middle, the reference DNA region (CHR4) and the reference sample (Normal) used in this dataset. Bottom, the confidence interval for determining a genomic instability, insertion, or deletion event. (**D**) Bar chart showing the output from Auto-qPCR program running the genomic instability model. Four different iPSC cell lines are indicated and compared to the control sample. Normalized signals for all four cell lines are in the confidence interval defined by the control sample.

## Results

### The Auto-qPCR program functions with the workflow of a qPCR experiment

A qPCR experiment includes multiple steps that can be divided into two categories: (1) sample preparation to conduct the qPCR reaction, and (2) data analysis, visually represented in the schematic in **Figure 1**. Nucleic acids are extracted from biological samples (RNA which is converted to cDNA for quantifying gene expression levels; or genomic DNA). Prior to performing qPCR *in vitro*, the user must generate the *in-silico* experimental layout using software that monitors the biochemical reaction. The user defines the experimental design (absolute or relative quantification), the method for detecting DNA synthesis (Taqman or SybrGreen) and the location of each sample within the plate. Finally, at the end of the qPCR process/cycle/program, the recorded data is exported and then would normally be analyzed manually. In our workflow, the data is exported from the PCR machine and saved as spreadsheet in the form of a txt or csv file **(Supplementary Figure S2)**. The file is then uploaded into the Auto-qPCR web app and the user enters their experimental settings.

Auto-qPCR will remove technical replicates by the selected criteria, normalize to an endogenous control, create a clean data table, and summary data table and graphs of all the results. If the user selects the statistical analysis, differential expression analyses will be performed on the designated groups. The program was designed for the most common uses of qPCR: detecting DNA fragment duplications or deletions, and quantifying gene expression levels according to the absolute or relative quantification models.

### Genomic instability

A relatively new application for qPCR detects small changes within the genome, from a deletion to a duplication of a DNA segment. DNA regions known to be highly susceptible to such events can be quantified using a genomic instability qPCR test. In induced pluripotent stem cell (iPSC) research, genomic instability tests are critical for quality control to screen for duplication/deletion events that can arise during reprogramming and prolonged cell passaging ^30, 31^. We performed a qPCR test for genomic stability, where for each cell line, the signal from each DNA region of interest was compared to the endogenous control region.

We uploaded the data into the Auto-qPCR web app and selected the genomic instability model **(Fig. 2B)**. The endogenous control used to normalize the data, was an amplicon of a region on chromosome 4 (CHR4), a location of the genome known not to contain any instabilities. As a reference sample, we used DNA known not to have any instabilities as the calibrator (Normal) **(Fig. 2A)**. The genomic instability model has two steps of normalization in its general formula. This formula and the variables used in the example calculation (**Fig. 2B and C**). First, the CT values from the control region (i.e., CHR4) for each cell line are subtracted from each region of interest. Next, the ΔCT from the Normal DNA control is subtracted from the ΔCT calculated for each cell line sample. Finally, the mean is calculated from the average of multiple technical replicates included with the plate design for each sample. Thus, the ΔΔCT values are expressed as “Relative Quantification” according to the following formula: RQ=2^-ΔΔCT^. If the sample has no abnormalities (deletions or duplications) the values obtained should be equal or close to 1, except for targets in the X chromosome in a male individual in which the ratio would be expected to be at 0.5. As the DNA used for PCR amplification may come from a mixed population of cells, where only some cells carry a deletion or duplication, we set an acceptable range of variation as 0.3 above and below the expected value of 1; DNA regions with RQ values between that 0.7 and 1.3 are considered normal. Values below 0.7 indicate a deletion and values above 1.3 indicate an insertion. For ease of analysis, we have included a column in the output file from the Auto-qPCR program that indicates normal, insertion or deletion **(Supplementary Table S6)**. We found that all seven chromosomal regions in the four cell lines tested were between 0.7 and 1.3 and we concluded that no duplications or deletions were present **(Fig. 2D and Supplementary Fig. S3B)**. Overall, we demonstrated how Auto-qPCR can be used to analyse the data from a genomic instability qPCR assay, and that the app effectively processed the data, creating a summary table and graph of the data.

### Absolute Quantification

For absolute quantification experiments, the quantities of RNA transcripts for a gene of interest and the endogenous controls are first estimated with a calibration curve **(Fig. 3A)** to provide a mathematical relationship between the CT values and the RNA concentration or quantity. The relationship is described by the equation CT=alog2[RNA] + b, where “a “is the slope and b is the Y-intercept **(Fig. 3C)** ^32^. The expression levels of the RNA molecule of interest are then given by the ratio of the estimated amount of RNA for a select transcript and the estimated amounts of endogenous controls **(Fig. 3C**). Consequently, the values given as “Normalized Expression Levels” depend on the levels of transcript within the biological material used to set the calibration curves. We used Auto-qPCR to compare the expression of three gene transcripts across six different cell lines at four different stages in the differentiation of neurons from iPSCs **(Fig. 3B and Supplementary Fig. S4)**. The calibration curve was made from a mix of the cDNAs generated from the reverse-transcribed RNA reactions from the four timepoints in the differentiation process and made of eight four-time serial dilutions to cover a linear relationship in a dynamic range from 1 to 16384-fold dilution **(Fig. 3A)**. Raw data was normalized with two endogenous controls (*ACTB and GAPDH*) **(Fig. 3D-H and Supplementary Fig. S4A)**. Auto-qPCR app provides several graphical representations of the normalized expression values. The means of technical replicates are provided for each gene **(Fig. 3D)**. Bar charts were generated for all gene and sample observations plotted together (grouped by gene **Fig. 3E** and by sample **Fig. 3G**), allowing for an overview of the data and visualization of the biological variation between cell lines at a given stage.

**Figure 3:**
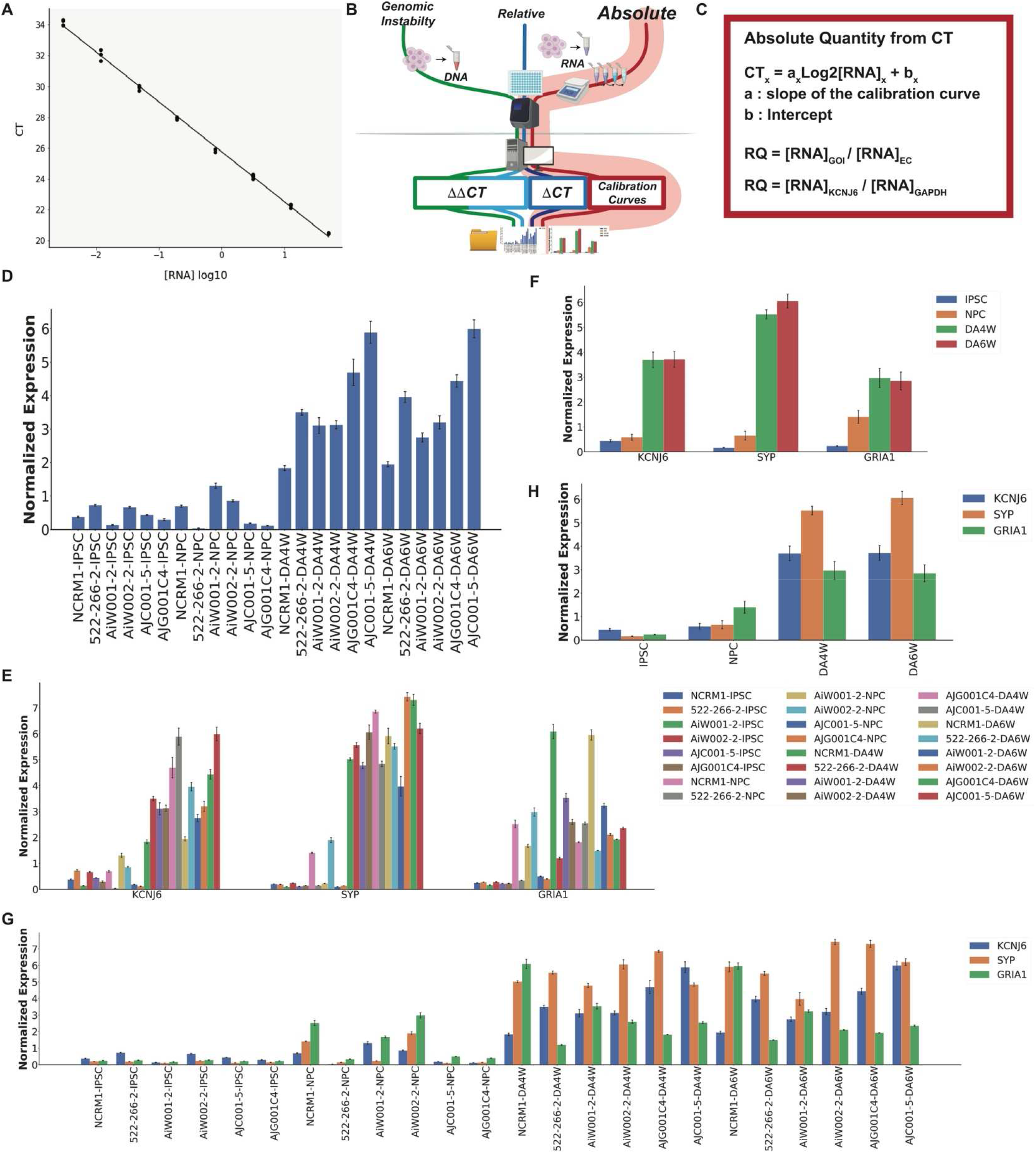
Auto-qPCR can process quantitative qPCR data using a standard curve to perform statistical analysis. Output of Auto-qPCR processing using the absolute model. (**A**) Illustration of a calibration curve displaying 8 serial dilution points of a four-fold dilution which covers cDNA quantities from 0.003053 to 50 ng and establishes the linear relationship between CT values (y-axis) and the log2[RNA]. (**B**) Schematic of PCR workflow showing the pipeline for the absolute quantification using a standard curve in red. (**C**) Formula used to process a real-time PCR experiment using an absolute quantification design. Top, general formula where the linear relation between the logarithm of RNA concentration and the CT value is provided by the calibration curve. The normalized quantification is expressed as a ratio between concentrations for the gene of interest and the endogenous control(s) estimated from their respective calibration curves. Bottom, the variables specific to this dataset are shown in the general formula. (**D**) Bar chart showing the output from Auto-qPCR program using the absolute model for the normalized expression of the gene *KCNJ6* for six cell lines at four different developmental stages (iPSC, induced pluripotent stem cells; NPC, Neural progenitor cells; DA4W, dopaminergic neurons at 4 weeks, DA6W: Dopaminergic neurons at 6 weeks). (**E**) and (**G**) Bar charts showing the average expression levels obtained from the three technical replicates for each cell line and time point for the three genes (S*YP, KCNJ6* and *GRIA1),* normalized with two housekeeping genes (*ACTB*: *beta-actin*, *GAPDH*). (**E**) Mean RNA expression grouped by genes on the x-axis, cell lines and time points are indicated in legend. (**G**) Mean RNA expression grouped by cell lines and time points; the gene transcripts quantified are indicated in the legend. (**F**) and (**H**) Bar charts showing the mean expression levels of S*YP, KCNJ6* and *GRIA1* for four developmental stages (n=6 cell lines). (**F**) Grouped by genes (x-axis), time points are indicated in the legend. (**H**) Grouped by time points (x-axis), the genes are indicated in the legend. One-way ANOVAs across differentiation stages for *KCNJ6*, *SYP* and *GRIA1* (p < 0.001, p < 0.001, p = 0.002).

We used the statistical module in Auto-qPCR to test for changes in gene expression over the different stages of neuronal differentiation; the different cell lines were considered as biological replicates **(Supplementary Fig. S5)**. As there are more than two groups, the Auto-qPCR software runs a one-way-repeated measures ANOVA for each gene. Two summary plots **(Fig. 3F and H)** and two statistical output tables were generated: one for the ANOVAs and one for the secondary measures **(Supplementary Tables S7 and S8)**. There was a significant effect of the differentiation stage on the expression of synaptic markers. The t-tests with false discovery rate (FDR) correction for pairwise comparisons of each stage showed that iPSCs have significantly less expression of each synaptic marker than DAN differentiated for 4 and 6 weeks **(Supplementary Table S8)**, indicating that the differentiation protocol is successful for all cell lines tested, with each iPSC differentiating into progenitors and ultimately DAN **(Supplementary Figure S5)**. We show that raw absolute qPCR data was effectively processed by Auto-qPCR, creating summary data, visualization and statistics for differential gene expression between conditions.

### Relative quantification

In addition to absolute quantification, the Auto-qPCR software also enables the processing of qPCR data obtained according to a relative quantification design. Contrary to absolute quantification, relative quantification does not require a calibration curve, and quantification (of transcripts) is based on the CT difference between a transcript of interest and one or more endogenous controls **(Fig. 4A)**. Relative qPCR is optimal for two kinds of comparisons: (1) detecting a difference in gene expression between two different conditions, and (2) detecting a difference between two transcripts within the same condition. Relative quantification can be expressed either as RQ=2^-ΔCT^, where samples are normalized to internal control(s), or RQ =2^-ΔΔCT^, where a given sample is considered as a calibrator for the unknown samples **(Fig. 4B and C)**.

**Figure 4.**
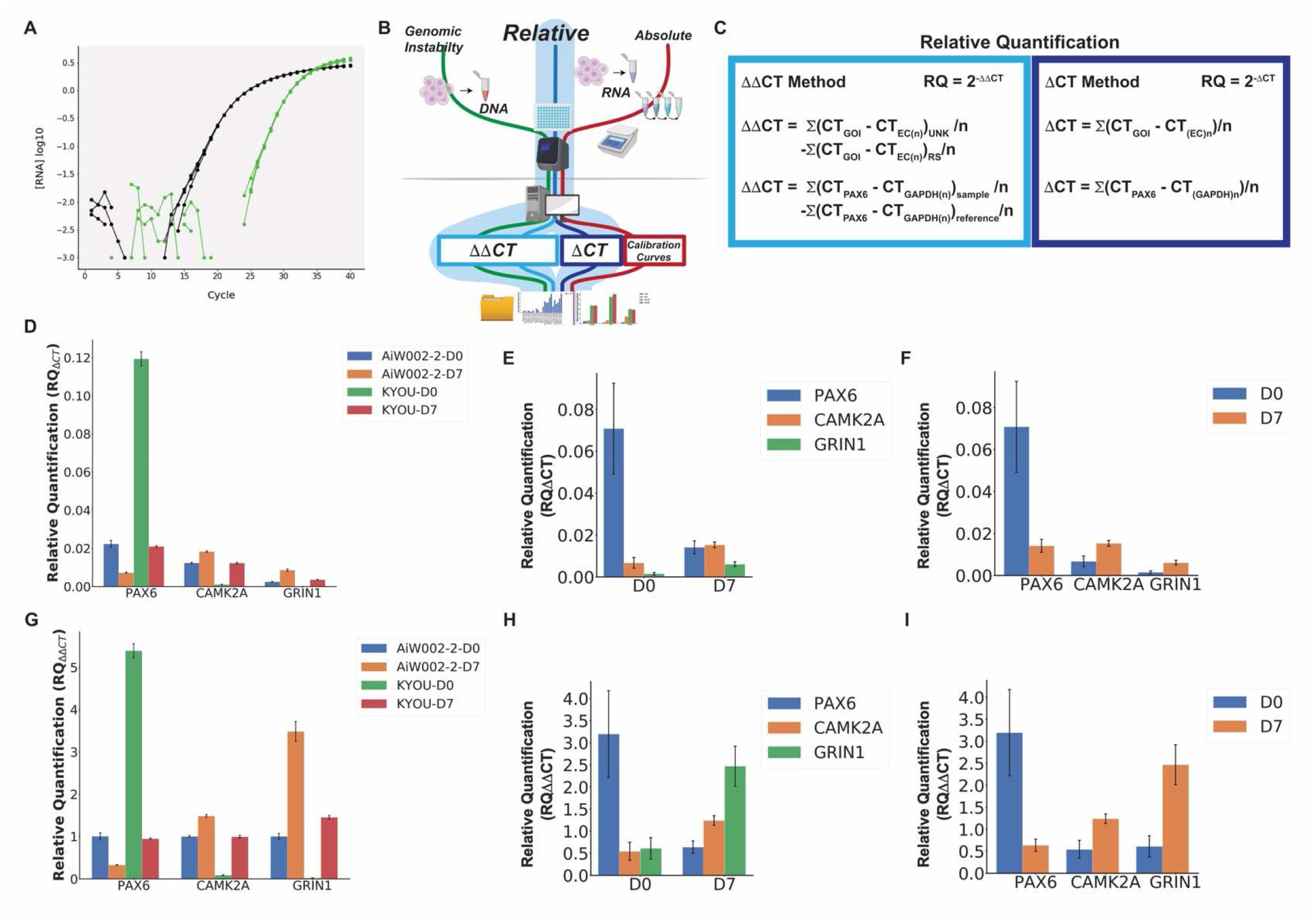
Auto-qPCR can process quantitative PCR data using two different relative models. Output of Auto- qPCR using the relative quantification with both the ΔCT and ΔΔCT models. (**A**) Amplification curves illustrating a difference of cycle threshold values (ΔCT) between a gene of interest and an endogenous control. (**B**) Schematic of PCR workflow showing the two methods to calculate relative RNA quantity, ΔCT in dark blue and ΔΔCT in light blue. (**C**) Formula used to perform a qPCR using relative quantification models, according the ΔCT (right), or the ΔΔCT methods (left). (**D-F**) Bar charts showing the output of the delta-CT model (RQ^ΔCT^). **G-I**) Bar charts showing the output from the ΔΔ-CT model (RQ^ΔΔCT^). (**D**) and (**G**) Mean normalized gene expression values from technical replicates for the genes *PAX6*, *CAMK2A* and *GRIN1* indicated on the x-axis for 2 cell lines at two stages of differentiation (D0: Neural progenitor cells, and D7: cortical neurons at 7 days of differentiation) as indicated. (**E**) and (**H**) Statistics output showing the mean gene expression from two cell lines at two stages of differentiation indicated, for the three genes indicated on the x-axis. (**F**) and (**I**) Statistics output showing the mean expression values for two cell lines at two time points on the x-axis and the three genes indicated. Differential expression between D0 and D7 is not significant (*PAX6* p = 0.40, *CAMK2A* p=0.18, *GRIN1* p=0.16), t-tests, n=2.

To illustrate the functions of the program, we compared the expression levels of two different control cell lines at two developmental stages, indicated as D0 (neural precursor cells) and D7 (7 days of differentiation into cortical neurons). We measured the expression levels of the progenitor marker *PAX6*, and two markers of neuronal differentiation (*GRIN1*and *CAMK2A) and* normalized to the housekeeping genes *ACTB* and *GAPDH.* We used the Auto-qPCR app to process the same data twice, for a direct comparison of the two distinct relative quantification options **(Supplementary Fig. S6)**. **Figure 4D** shows the mean expression from technical triplicates calculated by selecting the RQ=2^-ΔCT^. The ΔCT approach (not using a sample as calibrator) allows a comparison of the expression levels for the three different transcripts. We observed that relative to the endogenous controls, the D0 expression values for each transcript varied widely between the two cell lines tested. However, as expected for both cell lines, *PAX6* expression is higher at the D0 stage compared to D7. Conversely, both *GRIN1* and *CAMK2A* exhibited higher expression at the D7 stage compared to D0. Using the statistics module in the Auto-qPCR app, we compared the mean levels of each gene transcript at D0 and D7 using paired t-tests for each gene **(Fig. 4E and F)**. We found that although there were clear differences in expression, they were not significant between D0 and D7, likely a result of there only being two samples for each time point **(Supplementary Table S9 and Supplementary Fig. S6A and S7)**. Interestingly, we found that the *CAMK2A* RQΔCT was twice the level of *GRIN1* at D7 RQΔCT (**Fig. 4F**).

We next analysed this dataset with the RQΔΔCT model (indicated as ΔΔCT) in the web app **(Supplementary Fig. S6B)** where transcript levels are compared to both control gene expression (in this case *ACTB* and *GAPDH*) and a calibration sample; in this case we set one sample, AIW002-02-D0 arbitrarily as the reference sample **(Fig. 4G)**. Here we can easily compare expression in a test condition relative to a control condition by displaying the results as fold change in expression. All decreases are displayed as between 0 and 1 and all the increased expression levels are above 1 **(Fig. 4C)**. With the double normalization (RQΔΔCT), all values were expressed as a variation compared to the calibrator (AIW002-2-D0) as seen in Figures 4G-I. As in the RQΔCT model, the changes in gene expression from D0 to D7 were not significant **(Supplementary Table S10)**. Although the ratio of expression for a given gene in each cell line between DO and D7 remained unchanged, differential expression between genes can no longer be analysed. The RQΔΔCT shown in **Fig. 4H** showed that *PAX6* expression was higher at D0 than D7 and that *CAMK2a* and *GRIN1* expression were both higher at D7 than D0, as seen in **Fig. 4E** using the RQΔCT model. However, with the double normalization, the increase in *GRIN1* expression from D0 to D7 appears much larger than the increase in *CAMK2a* expression **(Fig. 4H and I)**, which was the opposite result from the single normalization model (RQΔCT) **(Fig. 4E and F)**. Our findings highlight the need to analyze data with attention to the biological question. Using only the RQΔΔCT analysis, one might mistakenly believe the increase in *GRIN1* expression is greater than that of *CAMK2a.* With Auto-qPCR we provide a quick easy option to process the exported qPCR data with two different relative models. We show the same gene expression ratios between the two time points, but different expression gene levels using the different relative quantitation models.

### Auto-qPCR produces the same results as manual processing of a previously published dataset

One of our objectives was to provide a tool for analyzing data from qPCR experiments generated with different qPCR machines. We reanalyzed a published dataset generated by the Gorwood lab ^29^, on a different machine (Opticon 2, Biorad). The original study measured gene expression in three sub cortical areas (subthalamic nucleus (STN), substantia nigra (SN) and globus pallidus (GP) of mice submitted to a place preference paradigm to cocaine ^29^. Manual processing shows a significant increase in *Nrxn3* expression in the cocaine-treated group compared to control, specifically in the GP **(Fig. 5A)**.

**Figure 5:**
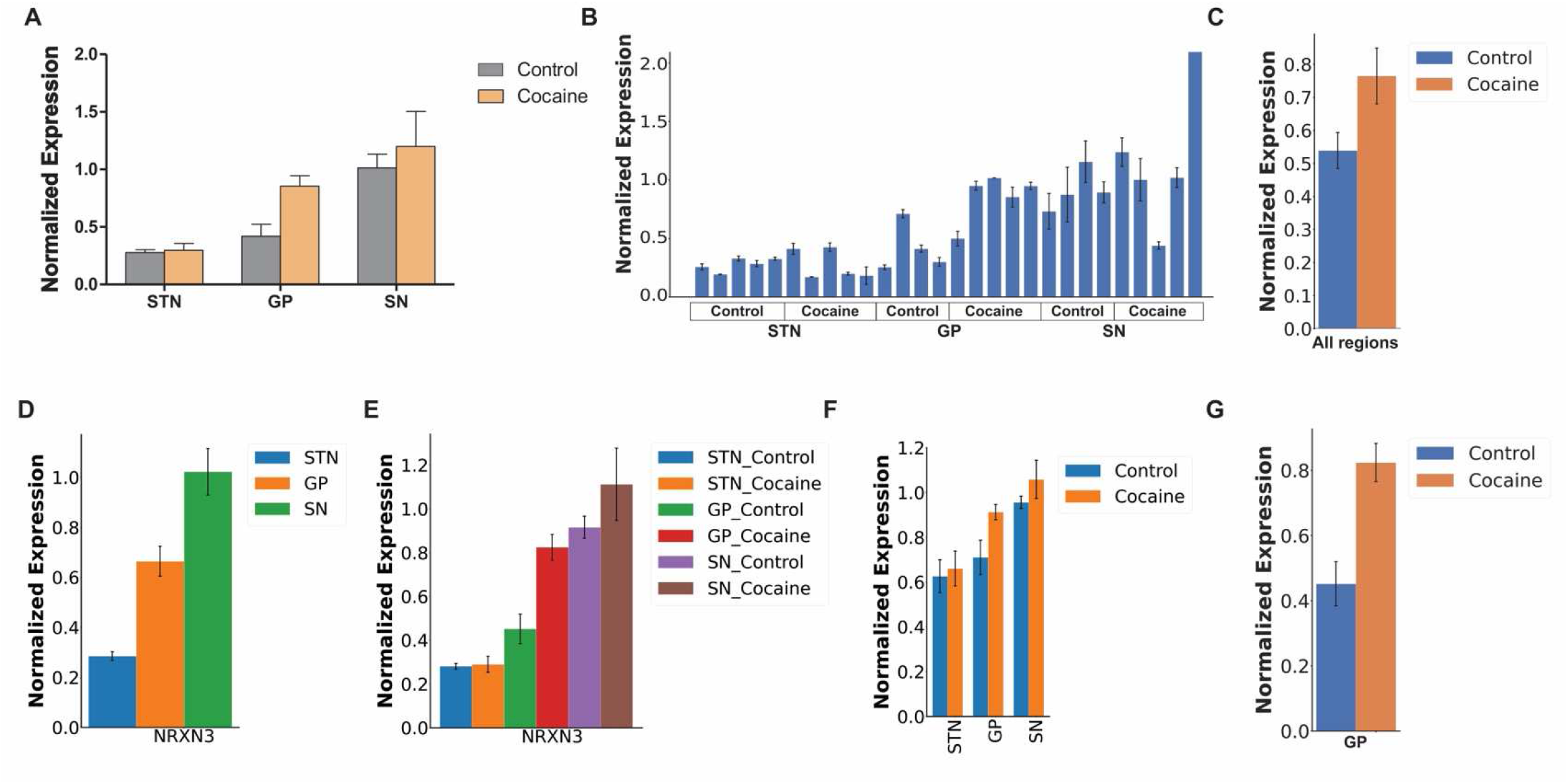
Auto-qPCR can process data from different thermocyclers and produce the same results as manual processing. (**A**) Bar chart showing the mean *Nrxn3* expression level normalized to *B2M* levels assessed with an absolute quantification design manually processed and plotted in Prism, grouped by brain regions (STN: subthalamic nucleus, GP: globus paladus, SN: substantia nigra) on the x-axis, with and without cocaine treatment. (**B**) Output of Auto-qPCR processing the same dataset. *Nrxn3* normalized expression levels from technical replicates for each biological sample. The treatment conditions are indicated below the x-axis. (**C**) Statistics output of Auto-qPCR program comparing cocaine and control groups. *Nrxn3* normalized expression levels in the combined brain regions. Expression is not significantly different, p=0.113, t-test, n=13. (**D**) Auto-qPCR statistical output showing mean *Nrxn3* expression combining treatments and comparing the three brain regions. One-way ANOVA shows significant effect of brain regions, FDR adjusted p < 0.001, n=9 for GP and SN, n= 10 STN. (**E**) Bar chart of Nrxn3 expression shown as six groups distinguished by brain region and treatment generated by Auto-qPCR program after a one-way ANOVA, p < 0.001, n=4 or 5. Posthoc analysis using multiple t-test with FDR correction comparing treatment at each brain region: SNT p=0.990, GP p=0.033, SN p=0.413. (**F**) Bar chart of *Nrxn3* average normalized by brain region (x-axis) and treatment, generated by Auto-qPCR program after a two-way ANOVA, brain region p < 0.001, treatment p= 0.2265, n=4 or 5. Posthoc analysis using multiple t-test with FDR correction comparing each brain region with and without cocaine: SNT p=0.0.998, GP p=0.053 and p-unadjusted = 0.017, SN p=0.619 (**G**) Bar chart of the average *Nrxn3* normalized expression levels in the GP compared between the two groups with a t-test (p = 0.0176).

We next processed the raw data using the Auto-qPCR web app absolute quantification pipeline and normalized to *B2M* **(Fig. 5B and Supplementary Figure S8A).** This summary data closely matched the manually calculated data **(Supplementary Table S11)**. The standard method of removing outliers from technical replicates is to remove the replicate most different from the mean, if the CT standard deviation (std) is above 0.3. Under ‘Options for removing technical replicates’ in the Auto-qPCR software the threshold can be adjusted. During manual analysis, each set of technical replicates is inspected when the std value is above 0.3, when one replicate is clearly different from the other two the divergent value will be removed. There are some instances in manual processing where no replicates are removed when the std is greater than 0.3, because the triplicate values are evenly distributed. Auto-qPCR has an option to account for this type of data when the user selects ‘preserve highly variable values’. With this option a replicate is only removed if the median is far from the mean. We processed the *Nrxn3* expression data with a range of std cut-off values to display the difference in outcomes and with or without preserving highly variable replicates **(Supplementary Table S11)**. We compared the variances generated by the differences between the expression values from manual treatment and from Auto-qPCR using a std cut-off of 0.3 with or without preserving highly variable replicates. We found that the preservation of highly variable option combined with a cut-off at 0.3 generate a 20% decrease in the variance between manual and automatic treatments **(Supplementary Table S12)** and preserved values falsely estimated as outliers by manual processing, which illustrates the subjectivity of the user with respect to the decision to retain or exclude a value based on criteria of divergence Our analysis suggests that applying two rules of data filtering provides a more systematic data analysis method and minimizes interindividual bias. Here we applied the standard cut-off of 0.3 and preserved highly variable replicates, appropriate for the highly variable and RNA level experimental samples we are analyzing.

Auto-qPCR also permits statistical groups to be designated in the sample name or in a specific group column, which can be added into the qPCR data during the plate set up or later in the results spreadsheet. To allow for statistical analysis of this data, we added a grouping column into the raw data files **(Supplementary Table S13)** and using the Auto-qPCR statistics module, we reanalysed the effect of drug treatment and brain regions on expression of *Nrxn3* across several parameters. We first compared the overall effect of cocaine on expression after pooling the three brain regions and found that although the expression of *Nrxn3* was increased across brain regions with cocaine treatment, there was no overall significant effect of drug treatment **(Fig. 5C, Supplementary Fig. S9A and Supplementary Table S14)**. Comparing the three brain regions while pooling together control and cocaine treatment showed a significant difference in expression across brain regions. Post-hoc analysis revealed *Nrxn3* expression in the STN was significantly lower than in the GP and SN **(Fig. 5D, Supplementary Fig. S10A and Supplementary Table S15).** When we considered each brain region with and without treatment as independent conditions, and individual mice as biological replicates and used a one-way ANOVA followed by post hoc tests using multiple t-test with a correction for multiple comparisons we find cocaine significantly increased *Nrxn3* expression specifically in the GP and not in the SN or STN **(Fig. 5E and Supplementary Table S16**). To apply the identical statistical treatment as originally presented, we performed a two-way ANOVA followed by a repeated measures t-tests with FDR correction on the interaction variable between treatment and brain region, using Auto-qPCR, and found the same results as the one-way ANOVA **(Fig. 5F, Supplementary Fig. S10B and Supplementary Table S17)** and a t-test of the GP alone **(Fig. 5G)**, all in agreement with the originally published results ^29^. Together the data shows that the Auto-qPCR software is capable of processing data generated by another machine and the results match those processed manually.

## Discussion

This paper presents Auto-qPCR, a new web app for qPCR analysis and provides examples of the functionalities of the software applied to qPCR experimental datasets generated from DNA (genomic instability assay), cDNA amplification, and RNA transcripts (absolute and relative quantification data). We have also summarized the computational bases of relative and absolute quantifications performed by Auto-qPCR, which is important for users to understand during experimental design. The Auto-qPCR web app also provides a statistical module that will be applicable to the majority of qPCR analysis experiments, and provides a correction across multiple tests, when more than two samples are compared, to mitigate against false positives. As not all experimental designs require differential analyses, the user can use Auto-qPCR without statistical analysis, calculating normalized RNA concentrations, and a summary table and graphs will be generated. Furthermore, the web app can be used with no installation or login requirements. We have created an easy-to-use program that is completely free and open source, able to process data from different qPCR machines and all common experimental designs, that will be advantageous for any lab performing qPCR experiments.

Given the importance of qPCR in molecular biology, other programs are available to perform many steps of the qPCR data treatment ^18, 21–23, 33^. The Q-PCR and PIPE-T programs were designed to treat and display qPCR data generated according to a relative quantification model ^23, 33^. SATQPCR is a web app that treats qPCR data using the relative quantification model and performs differential analyses. However, it does not take the exported results files directly from the qPCR data and requires manually preformatting of the data before analysis ^22^. Finally, ELIMU-MDx is a web-based interface conceived to collect specific information regarding qPCR assays for diagnostic purposes. EILMU-MDx functions as a data management system, processes qPCR data generated using the absolute quantification method and requires an account and login information ^21^.

Reviewing different software published to serve similar purposes highlights the unique characteristics of Auto qPCR, as no other web app combines all the features we have included in our software. First as a web app, Auto-qPCR does not require installation or a user login and can be accessed from any device connected to internet. We also provide the option for users to install the program onto their computer if they want to work on their analysis off-line. Second, data processed by Auto-qPCR does not require any preformatting of the results file to be performed manually. Instead, once the qPCR experiment is complete, our program takes the csv or txt export file directly from the thermocycler so there is no copy/paste or formatting step to be done by the user. Third, Auto-qPCR can manage the data from multiple separate absolute files at once, as well as batch process multiple results files from a relative quantification. The program creates a clean data set (with all technical replicates) and a summary data table. Fourth, unlike the other software mentioned above, Auto-qPCR includes three different models, conceived to support qPCR data generated from absolute and two methods of relative quantification designs. No other program provides the option of choosing between the two relative quantification methods. Fifth, we provide normalization to multiple reference genes and calculate the mean normalized value for each replicate, and not the sample mean, an important feature implemented in relatively few other programs. This avoids the RNA quantity value being influenced by extreme values. Sixth, we extend the use of the program to suit qPCR data from DNA quantification. Finally, we provide an extensive statistics module for calculating differential gene expression that requires no additional input files. Options are included for experimental designs that include two or more sample comparisons (t-test, one- and two-way ANOVA and the equivalent non-parametric tests) and automatically generates bar charts for data visualization and summary tables with the statistical results. In summary, we have created a unique, easy to use qPCR analysis program that can benefit any researcher or lab that needs to analyze qPCR data on a regular basis, by saving time, avoiding errors and generating reproducible, figure-ready plots.

Auto-qPCR provides users the option for relative quantification by two methods: expression relative to endogenous control genes only (ΔCT method) or relative to endogenous genes and also normalized to a control condition (ΔΔCT method). Although the ΔΔCT method is considered the gold standard to express, in one number, the variation in gene expression between two conditions and the amplitude of that change in expression ^34^, it does not account for inter gene expression variation within the control condition ^35^. The differences between quantifying relative expression with or without a control condition used as a calibrator, are clearly demonstrated above **(Fig. 4)**. Expression levels of *GRIN1* and *CAMK2a* calculated with either relative quantification model were increased at seven days of differentiation (D7) compared to day zero (DO). However, we also found that *GRIN1* and *CAMK2A* had different levels in the baseline condition (ΔCT), thus we observe that information is lost when using a ΔΔCT normalization. For relative quantification using a ΔΔCT normalization we measured a fold change of variation compared to a control condition for a given gene ^36^, but information about differences of expression between two genes in control condition were not observed **(Fig. 4F)**. We have provided both the gold standard method of relative quantification and a method to calculate gene expression without a reference sample, to allow users to quickly determine expression changes without losing information about the level of expression in control conditions.

Reprocessing the external dataset highlighted two main advantages of treating qPCR dataset with a program. First, manual analysis of qPCR data is time consuming. Second, comparing both data treatments (manual and program-assisted) has shown that one important source of variation between results of manual analysis is the inconsistent rules used for data exclusion. Although removing one outlier from technical replicates, in the vast majority of cases, improves the CT standard deviation (std) by decreasing it under the commonly accepted threshold of 0.3, in many cases researchers decide to keep a technical replicate even if the CT-std value is above 0.3. These judgement calls frequently occur when transcripts have low expression levels and the high variance between technical replicates does not permit a decision based on the adjustment of the CT std. To account for these situations, we incorporated a second rule for data inclusion/exclusion based on the distance between the arithmetic mean and the median value of technical replicates to determine the most acceptable set of technical replicates. Applying such an algorithm to the user’s judgement removes variability and potential bias in the resulting normalized gene expression levels. We were able to reprocess external data using Auto-qPCR and acquired the same summary output, reaching the same conclusions as the initial study. We showed that Auto-qPCR can process data from different PCR machines and matched the expected outcome from manual processing without the risk of bias or errors. Using a double rule for data inclusion/exclusion for highly variable signal between technical replicates, the program provides a unique treatment that will considerably reduce the risk of variability and mistakes generated by and between users during manual data processing.

The Auto-qPCR program has some limitations and many other potential uses not included in this manuscript. Although the program is able of computing data from independent qPCR plates in singleplex (where each plate has a different amplicon), Auto-qPCR has not been adjusted yet to manage duplex qPCR (with one endogenous control and one transcript of interest quantified in the same well). Auto-qPCR has also not been equipped yet to process an inter-plate calibrator, required to cover a sample size of more than one plate, in absolute quantification mode experimental designs. Finally, as most of the primer sets for gene expression are now predesigned and eventually pretested by companies taking in consideration optimal efficiencies of amplification, correction factors for efficiencies have not been added into the Auto-qPCR algorithms. Despite these caveats, we propose that Auto-qPCR could be employed in a variety of molecular biology protocols. Auto qPCR is capable of analyzing data from a chromatin immunoprecipitation experiment followed by specific DNA amplification ^15^. The analyses could be performed using either the absolute or the relative quantification models. The absolute quantification method would permit testing primer efficiency through the calibration curve ^37^, and the DNA target amplification would be normalized to an unbound DNA as previously described ^38, 39^. Alternatively, the level of DNA/protein interaction can be estimated using the relative quantification models with one or several regions, known to be unbound by a protein of interest, as endogenous control(s) (ΔCT mode) and with a biological condition as a calibrator (ΔΔCT mode). Auto-qPCR is flexible enough to let the user choosing the most appropriate model to use, based on the information available on the DNA regions to amplify and analyze.

The Auto-qPCR program was conceived to treat, analyze, and display qPCR data generated using either relative or absolute quantification designs, while limiting errors related to manual processing. Data processing tools can’t replace or supplement appropriate experimental design and statistical power. The conditions included with the design and interpretation of the results still remain in the user’s hand. We have provided a tool that will provide easy, reproducible analysis without user errors for unlimited samples. Although, we cannot computationally remove the need for replication and controls, analysis time will no longer be a limitation. Auto-qPCR permits researchers to conduct studies with larger experimental designs while minimizing the risk of mistakes during the data analysis.

## Acknowledgements

T.M.D. received funding through the McGill Healthy Brains for Healthy Lives (HBHL) initiative, the CQDM FACS program, the Alain and Sandra Bouchard Foundation, the Ellen Foundation and the Mowfaghian Foundation.

T.M.D is supported by a project grant from CIHR (PJT – 169095). R.A.T was funded by a Healthy Brains for Healthy Lives Fellowship.

Thanks to Ivan Castanon Niconoff for helping create and set up the virtual machine used to host the Auto-qPCR web app. Thanks to Maria José Castellanos Montiel, Vincent Soubannier and Nguyen-Vi Mohamed, for testing the web app.

## Author Contributions

G.M. and R.A.T. conceptualized the program. I.D., G.G., E.C. and R.A.T. wrote and tested the program. R.A.T. managed the program development and GitHub repository and ran all the analysis using the webapp. G.G. built the graphical user interface and website. T.J.P.S. transferred the website to run online through a virtual machine. G.M. generated the qPCR data used to test the absolute and relative quantification models of Auto-qPCR program. C.X.Q.C., N.A. and A.N.J. extracted DNA and performed the PCR used to improve the pipeline related to the genomic instability model of Auto-qPCR program. S.K., N.R. and P.G. generated the external data set used for Fig. 5. R.A.T., I.D. and G.M. made the figures. G.M., R.A.T., L.K.B. and T.M.D wrote the manuscript.

## Supplementary Figure Legends

**Supplementary Figure S1.**
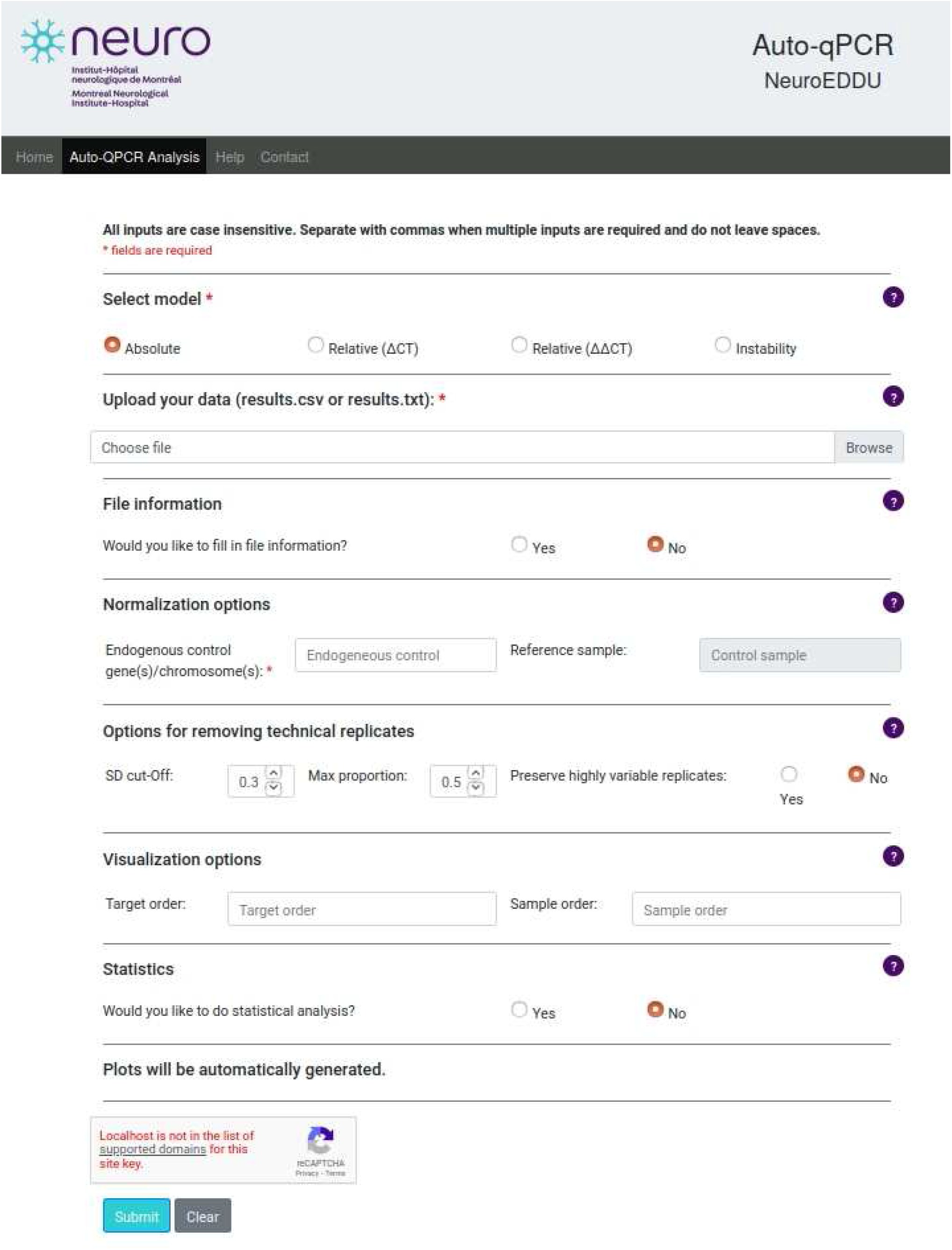
**Screen shot of the interface of Auto-qPCR.**

**Supplementary Figure S2.**
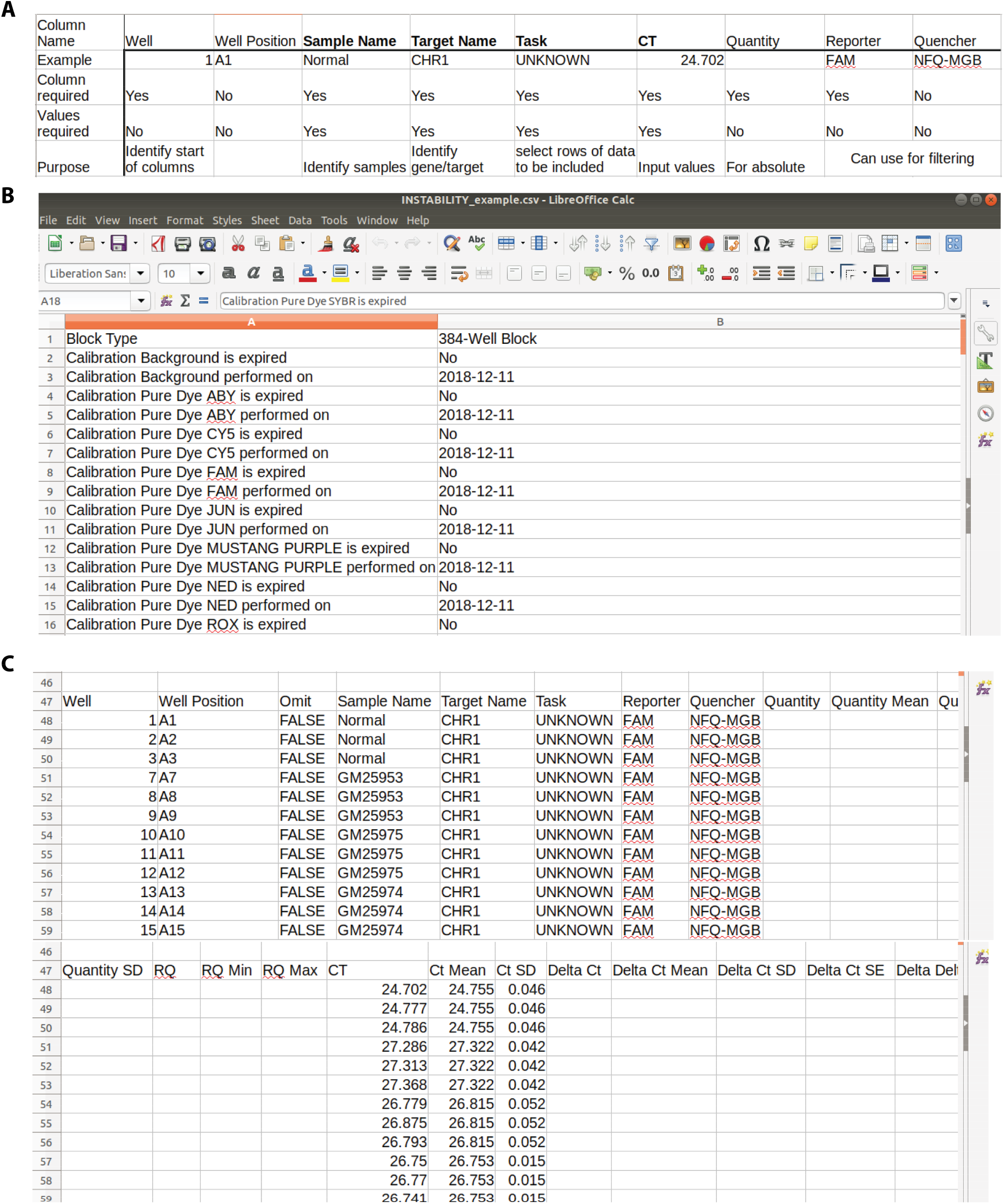
Examples of results spreadsheet files to use as input for Auto-qPCR. (**A**) Spreadsheet with column names needed. (**B**) Screen shot of the top of the csv saved from the results sheet of the exported excel file. (**C**) Screen shot of the column names in the save results file that will be read into Auto-qPCR.

**Supplementary Figure S3.**
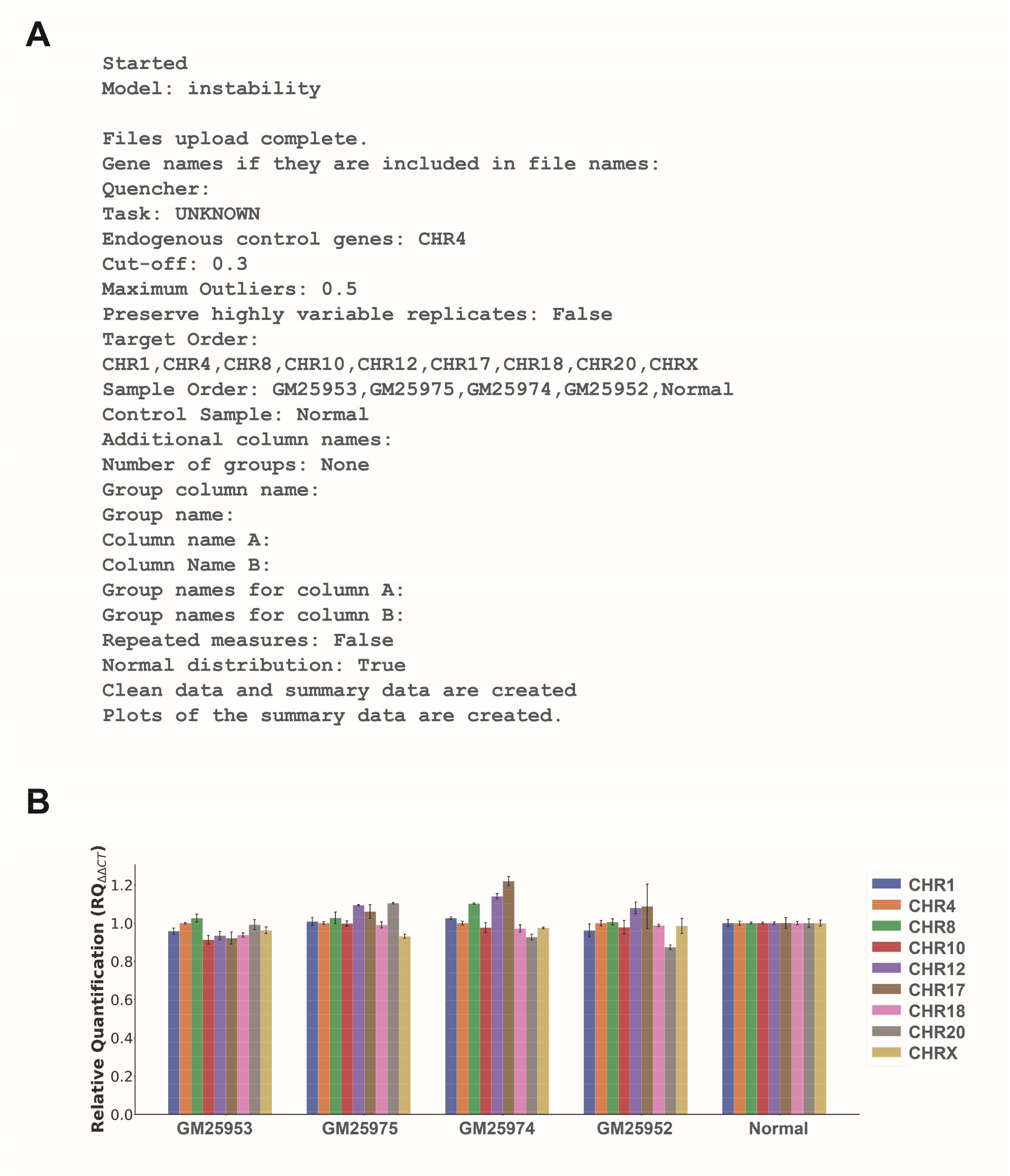
Example output from Auto-qPCR using the genomic instability model. (**A**) The Log.txt output from the file generated by Auto-qPCR. The file lists the steps completed by the program and the inputs from the web interface. This example is from the genomic instability analysis. The selection for statistical analysis is also shown in the text file. Using the log file, the exact analysis can be repeated because all the settings are recorded. (**B**) Bar chart showing an alternative visualization for the genomic instability assay where the data is grouped by cell lines on the x-axis and colours indicated in the legend represent the regions of chromosomes tested.

**Supplementary Figure S4.**
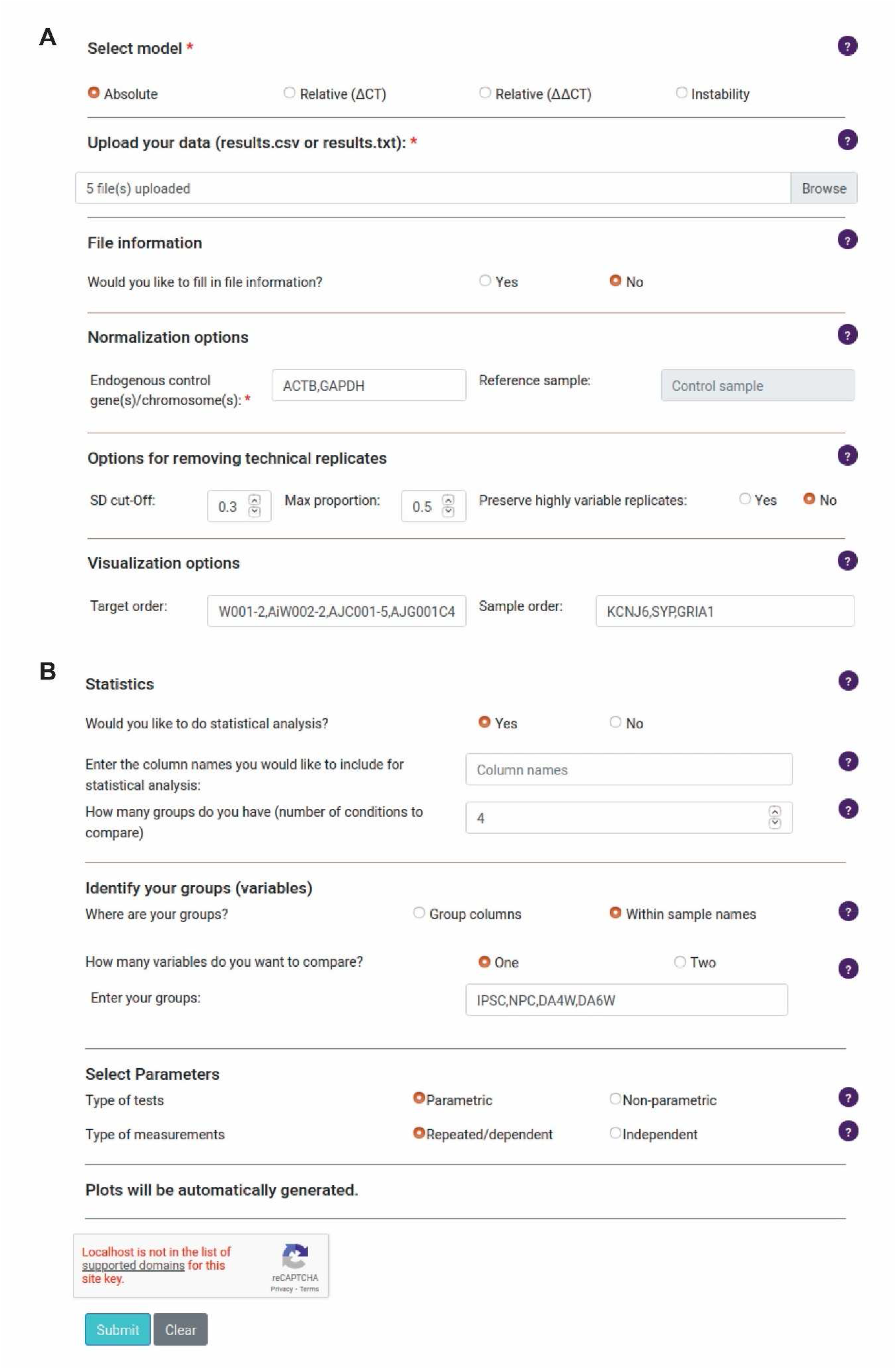
Screen shots of options entered into Auto-qPCR web app to analyze the example data for the absolute model in. Fig. 3. (**A**) Options to produce the summary data. (**B**) Statistics options.

**Supplementary Figure S5.**
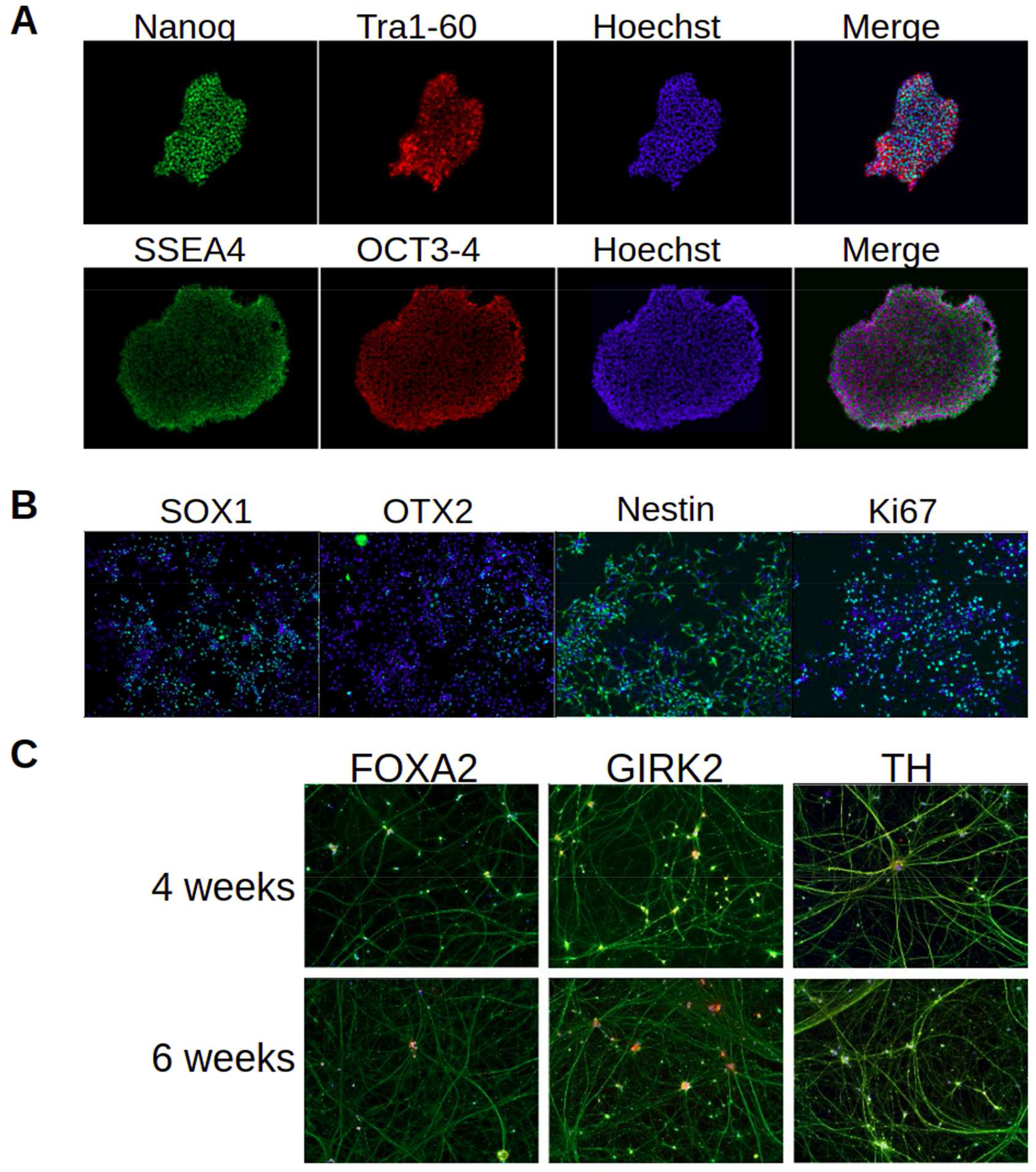
Example images of AJG001-C4 at four stages of development (iPSCs, NPCs, as well as 4 and 6 week DANs). (**A**) iPSCs stained for pluripotency markers (Nanog, Tra1-60, SSEA4, OCT3-4 as indicated), together with Hoechst and shown as merged images on the right. (**B**) Neural precursor cells (NPCs) expressing dopaminergic lineage (SOX1 and OTX2), proliferation (Ki67) and neural progenitors (Nestin) markers. (**C**) Dopaminergic neurons after 4 and 6 weeks of differentiation stained with neuronal marker Tuj1 in all images and dopaminergic markers FOXA2, GIRK2 and TH as indicated.

**Supplementary Figure S6.**
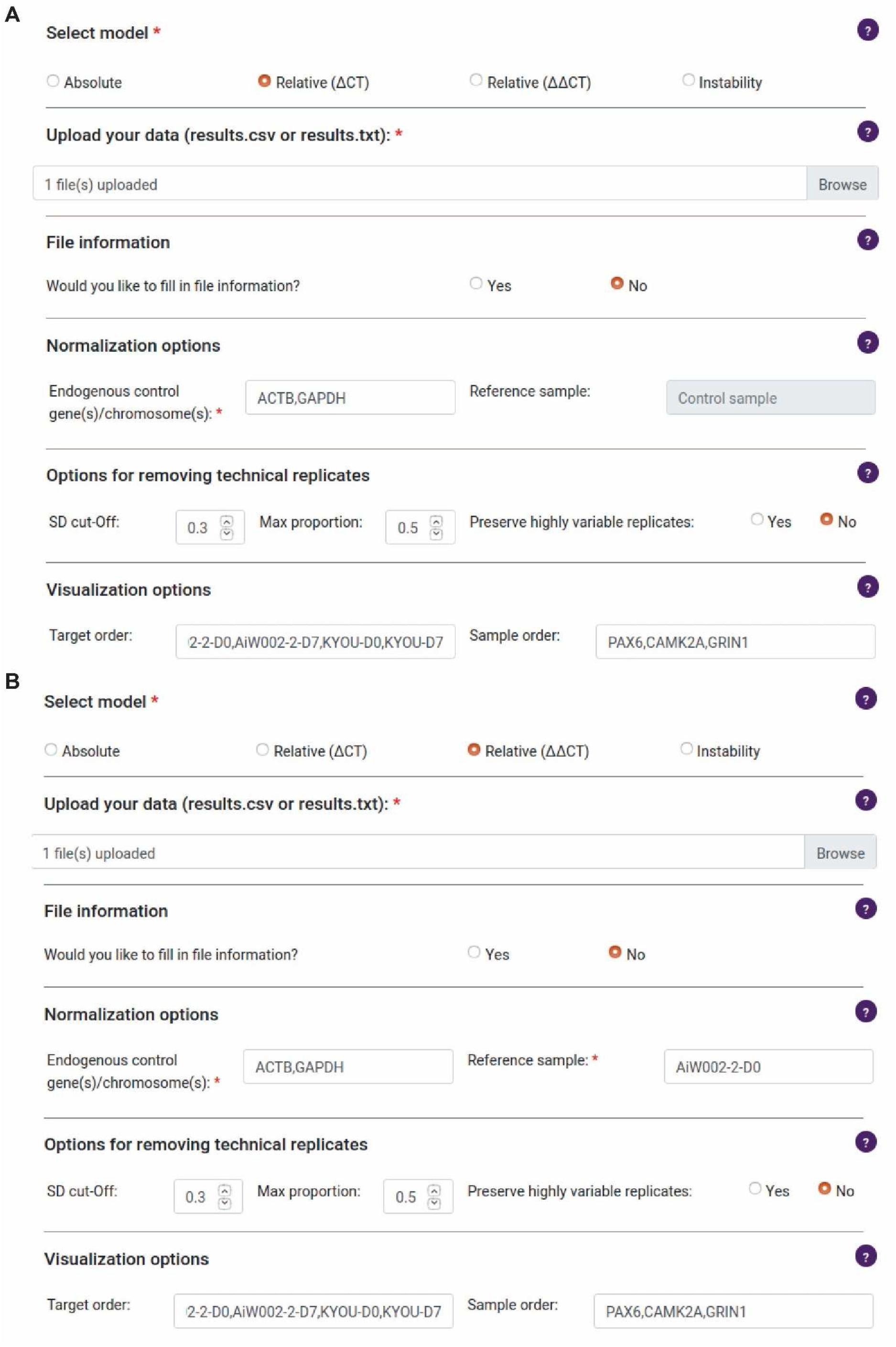
Screen shots of options entered into Auto-qPCR web app to analyze the example data for the relative models in. Fig. 4. (**A**) Options to produce the summary data using the relative ΔCT method, where values are normalized to the endogenous controls (ACTB and GAPDH). (**B**) Options to produce the summary data using the relative ΔΔCT method, where expression values are normalized both the endogenous controls and the reference sample.

**Supplementary Figure S7.**
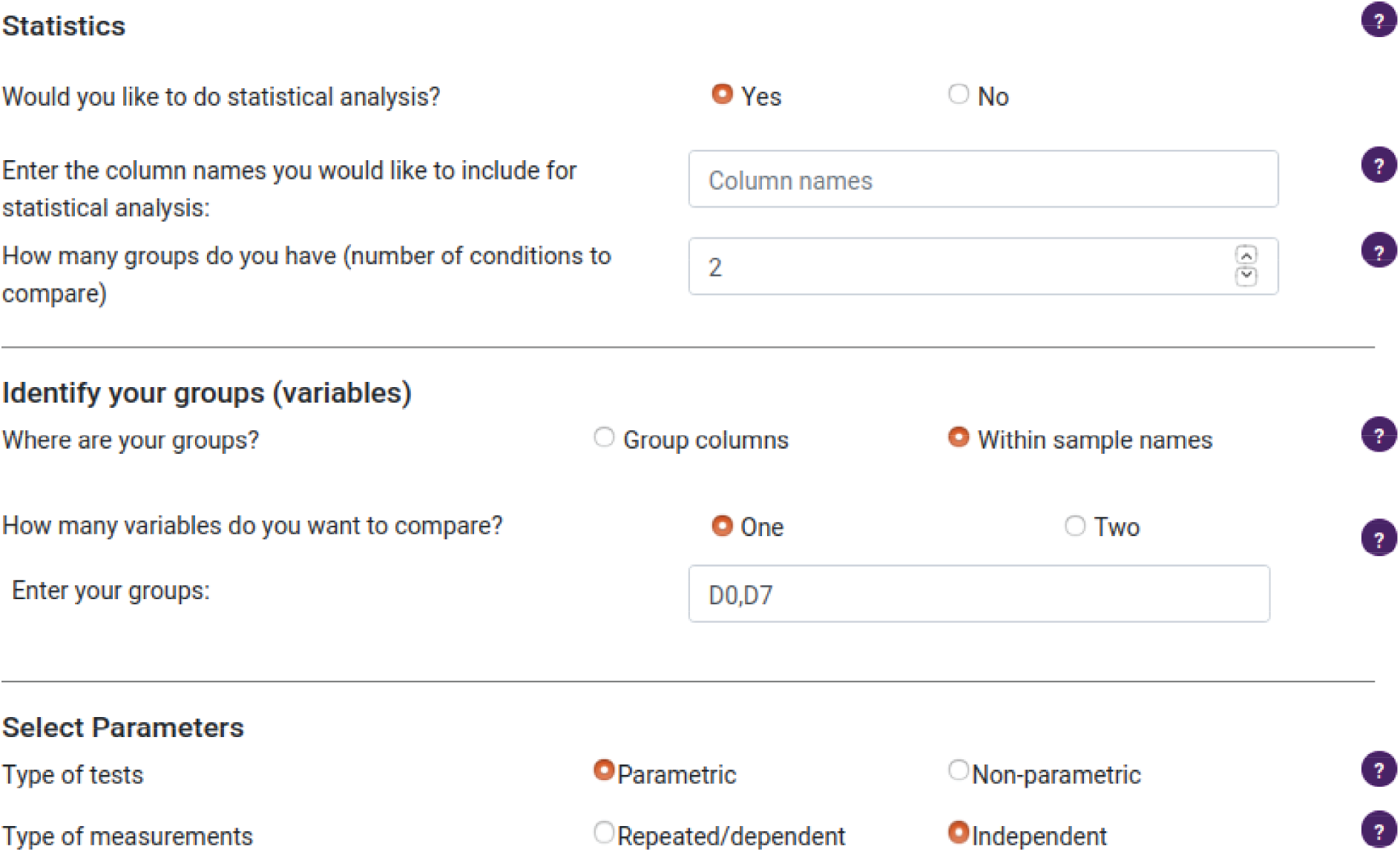
Screen shot of options entered into Auto-qPCR web app to for statistical analysis in. Fig. 4 **using relative models.** Statistics options used, the selections are the same for both the ΔCT and the ΔΔCT normalization methods.

**Supplementary Figure S8.**
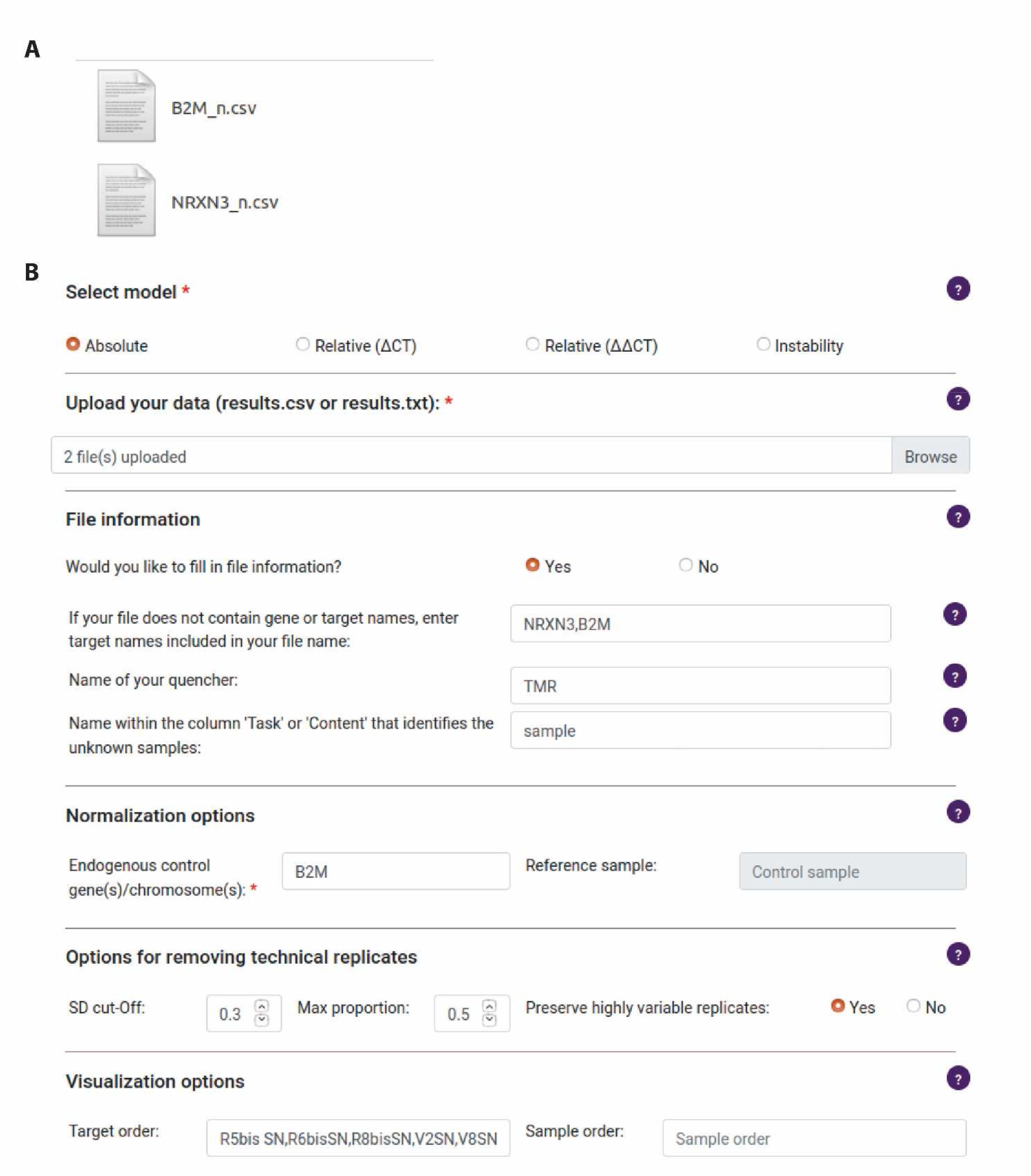
Screen shot of options entered into Auto-qPCR web app to for analysis of the input used for the absolute quantification to reprocess data from the Opticon 2 Biorad thermocycler. (**A**) Screen shot of file names that contain the endogenous control and the gene to be analyzed. (**B**) Screen shot of Auto-qPCR with the file names and entered under file information. All the options entered are to create the summary data used in Fig. 5.

**Supplementary Figure S9.**
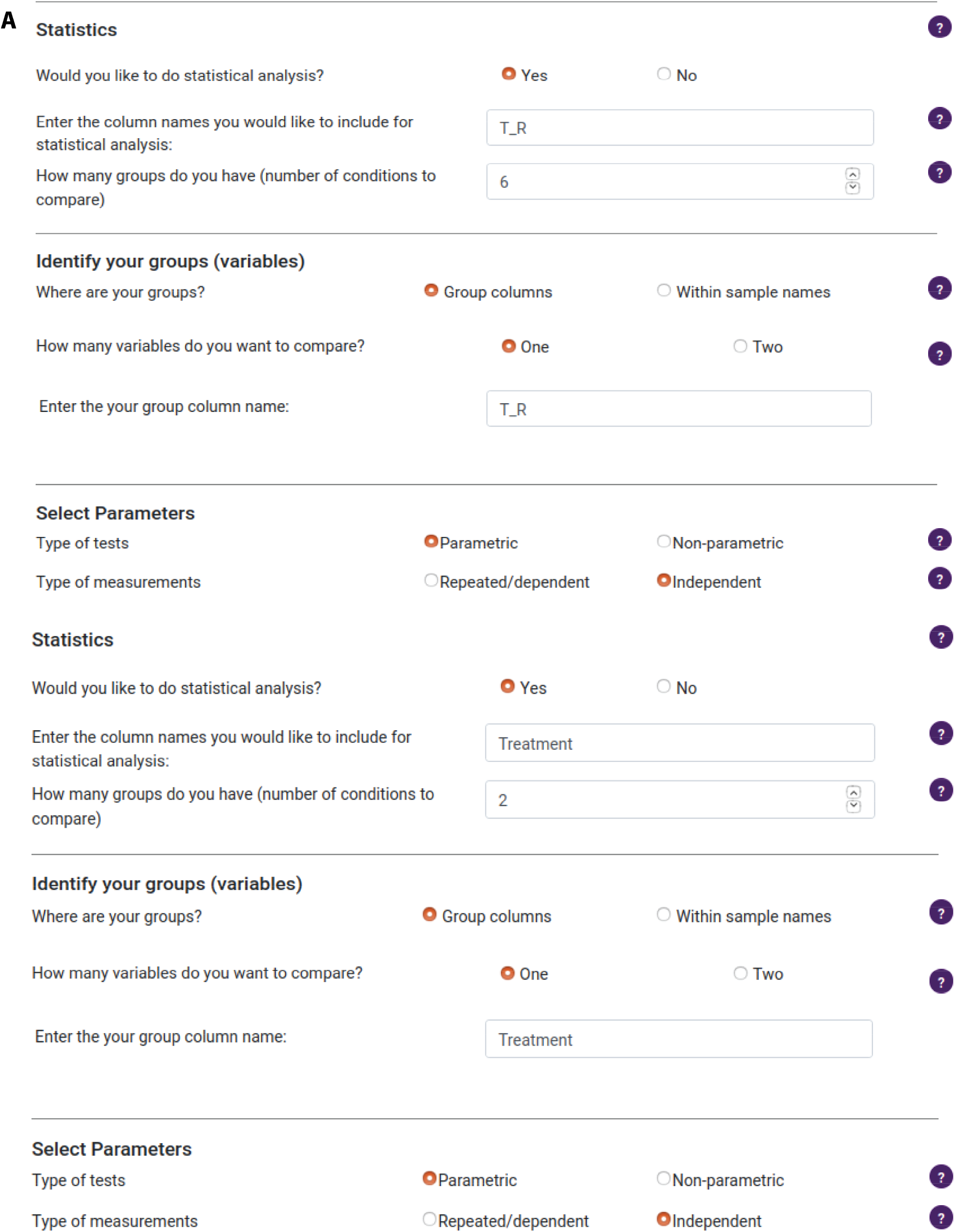
Screen shot of options entered for statistical analysis into Auto-qPCR web app for the absolute quantification to reprocess data from the Opticon 2 Biorad thermocycler. (**A**) Statistics options to compare brain regions and treatment combined to create 6 groups, a one-way ANOVA will be performed. (**B**) Statistic options to compare treatment and control (the brain regions are treated as one group), a t-test will be performed.

**Supplementary Figure S10.**
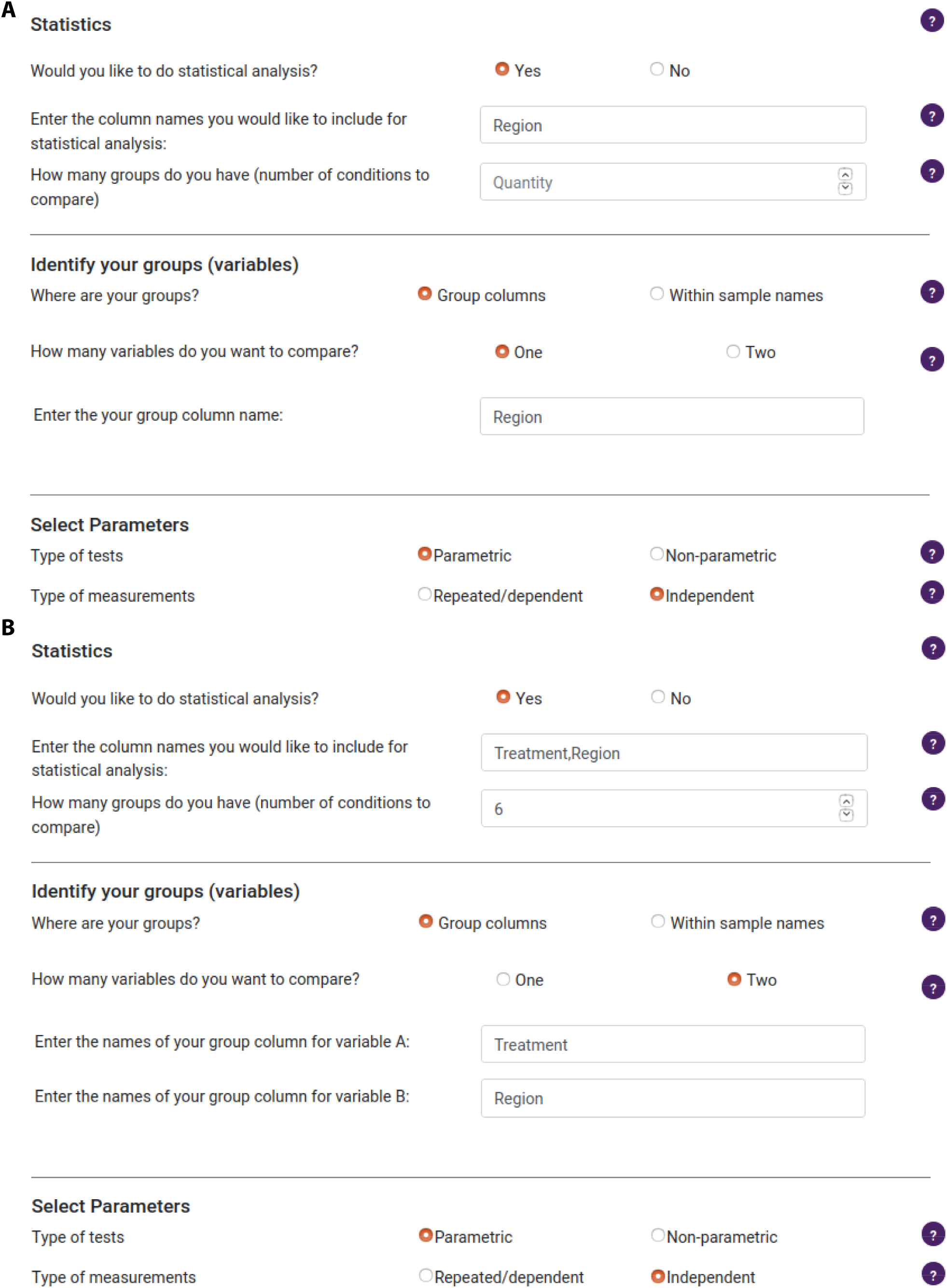
Screen shot of options entered for statistical analysis into Auto-qPCR web app for the absolute quantification to reprocess data from the Opticon 2 Biorad thermocycler. (**A**) Statistics options to compare brain regions with control vs. cocaine treated as one group, a one-way ANOVA will be performed with three brain regions as groups. (**B**) Statistic options for the two-way ANOVA where interaction between treatment and control is tested, the two variables are treatment and region.

## Supplemental Tables

**Table S1:** Overview of cell lines: Human-derived induced pluripotent stem cells used.

| Cell line | Donor Age | Sex | Cell Type | Reprogramming Method |
| --- | --- | --- | --- | --- |
| GM25952 | 10 | F | Fibroblast | Episomal |
| GM25953 | 43 | F | Fibroblast | Episomal |
| GM25974 | 7 | F | Fibroblast | Episomal |
| GM25975 | 37 | F | Fibroblast | Episomal |
| 522-2666-2 | NA | NA | Lymphocytes | Retrovirus |
| AIW001-2 | 48 | F | PBMCs | Retrovirus |
| AIW002-2 | 37 | M | PBMCs | Retrovirus |
| NCRM1 | NA | M | Cord Blood | Episomal |
| AJG001-C4 | 37 | M | PBMCs | Episomal |
| AJC001-5 | 37 | M | Fibroblast | Retrovirus |
| KYOU-DRX0190B | 36 | F | Fibroblast | Retrovirus |

**Table S2:** Taqman primers/probe sets. The primer/probe sets listed were used to generate the data presented in Figures 3 and 4 and test the absolute and relative quantification models to assess gene expression levels by Auto-qPCR web app. The primer/probe sets were selected from the assays available on the Thermo Fisher Scientific web site and chosen to cover the most important number of alternative transcripts for a given gene. With the exception of the assay for GAPDH, the amplicons overlap two exons, avoiding amplification of genomic DNA that could remain from incomplete DNAse digestion. The refseq sequence used for designing the primer/probe set assay is shown.

| **Gene Symbol** | **Gene Name** | **Location** | **Assay Reference** | **Exon Boundaries** | **Reference Accession** |
| --- | --- | --- | --- | --- | --- |
| *ACTB* | Actin beta | 7p22.1 | Hs01060665_g1 | 2-3 | NM_001101 |
| *GAPDH* | Glyceraldehyde-3-phosphate dehydrogenase | 12p13.31 | Hs02786624_g1 | 7 | NM_001256799 |
| *KCNJ6* | Potassium voltage-gated channel subfamily J member 6 | 21q22.13 | Hs01040524_m1 | 3-4 | NM_002240 |
| *SYP* | Synaptophysin | Xp11.23 | Hs00300531_m1 | 3-4 | NM_003179 |
| *CAMK2A* | Calcium/calmodulin-dependent protein kinase II | 5q32 | Hs00947041_m1 | 17-18 | NM_015981 |
| *PAX6* | Paired box 6 | 11p13 | Hs01088114_m1 | 7-8 | NM_000280 |
| *GRIN1* | Glutamate ionotropic receptor NMDA type subunit 1 | 9q34.3 | Hs00609557_m1 | 1-2 | NM_000832 |

**Table S3:** Contents and file structure of Python scripts. The file structure will be maintained if the Auto-qPCR program is downloaded from GitHub and run locally. These files will be found inside the ‘website’ folder if the GitHub repo is pulled or the zip file is downloaded. Folder Name indicates the parent folder and the subfolder containing the program files. File name indicates the file name for each Python script and Function indicates what processes are performed by each script.

| Folder Name | File name | Function |
| --- | --- | --- |
| Auto-qPCR | <u>main.py</u> | calls app |
| application | <u>AUTOqPCR.py</u> | inputs data |
|  |  | inputs conditions |
|  |  | removes outliers |
|  |  | calls model |
|  | <u>absolute.py</u> | runs normalization for absolute model |
|  | <u>relative.py</u> | runs relative quantification with delta-CT normalization |
|  | <u>stability.py</u> | runs relative quantification with delta-delta-CT normalization and genomic instability test |
|  | <u>plot.py</u> | creates all graphs |
|  | <u>statistics.py</u> | runs all statistics |
|  | <u>regex_rename.py</u> | function to allow flexible naming |
| application/template | all html interface files | creates the web form |

**Table S4:** List of all the user inputs for the Auto-qPCR program and purpose of the expected user inputs. Section indicates the spot in the web app where the input box is located. User Input indicates the input box or options as they appear in the web app. Selections and Values indicates possible options for the user to select and the purpose of the input.

| Section | User Input | Selections and Values |
| --- | --- | --- |
| Main | Select model | Choose the analysis model to run |
|  | Upload your data | Select your csv files |
|  | File information | Choose yes if your file doesn't contain gene names or you want to filter out data from a second probe. |
| Options | Endogenous control | Genes/targets for normalization |
|  | Cut-off | The threshold for which the standard deviation is above and outliers from technical replicates will be removed. Default = 0.3 |
|  | Max Outliers | The proportion of replicates that can be removed. Default = 0.5. With 0.5, if there are 3 replicates, only 1 can be removed |
|  | Preserve highly variable replicates | If set to yes, a second condition is added before a replicate is removed. The difference between the mean and median must be greater than 10 % of the mean |
| | Calibrator/reference sample | This is the gene/target that is the second normalization in the $\Delta\Delta CT$ model |
| Visualization Options | Target order | Genes are entered in the order they will appear on the graph |
|  | Sample order | Sample names are entered in the order they will appear on the graph |
|  | Columns for statistics | If a group column is present in the raw data, it must be indicated here to be available for the statistics |

**Table S5:** Description of the statistical tests using each possible selection criteria. The number of groups to compare, ‘#G’ indicates the number of conditions to compare with the variables. The number of variables, ‘#Var’ indicates the number of experimental conditions to compare. The distribution of the data determines if a parametric test will be used, for normally distributed data, or a non-parametric test will be used by the software. ‘Measure’ indicates if the data was collected on independent samples or on the same samples at different time points. ‘Test’ indicates the name of the test used by the software based on the user’s sections from the other four criteria. Auto-qPCR always uses the same post-hoc test except when only two groups are being compared and no post-hoc test is performed.

| # G | # V | Distribution | Measure | Test | Posthoc |
| --- | --- | --- | --- | --- | --- |
| 2 | 1 | parametric (normal) | Independent | student t-test two tailed, un-paired | none |
| 2 | 1 | parametric (normal) | Repeated measures (dependent) | student t-test two tailed, paired | none |
| 2 | 1 | non-parametric | Independent | Wilcoxon test | none |
| 2 | 1 | non-parametric | Repeated measures (dependent) | Mann-Whitney U test | none |
| > 2 | 1 | parametric (normal) | Independent | one-way ANOVA | pairwise t-tests with FDR correction |
| > 2 | 1 | parametric (normal) | Repeated measures (dependent) | one-way ANOVA | pairwise t-tests with FDR correction |
| > 2 | 1 | non-parametric | Independent | Kruskal-Wallis test | pairwise t-tests with FDR correction |
| > 2 | 1 | non-parametric | Repeated measures (dependent) | Friedman test | pairwise t-tests with FDR correction |
| > 2 | 2 | parametric (normal) | Independent | two-way ANOVA | pairwise t-tests with FDR correction |
| > 2 | 2 | parametric (normal) | Repeated measures (dependent) | two-way ANOVA | pairwise t-tests with FDR correction, for conditions 1,2 and the interaction |

**Table S6:** Results of Auto-qPCR summary output found in summary_data.csv. The DNA region is indicated in Target Name, cell lines are indicated in Sample Name, Indel indicates if there is a duplication or deletion event calculated by the web app, Rep is the number of technical replicates included for analysis, RQ is the relative quantification, Std is the standard deviation and SEM is the standard error of the mean. RQ values from the technical replicates.

| Target Name | Sample Name | Indel | Rep | RQ | Std | SEM |
| --- | --- | --- | --- | --- | --- | --- |
| CHR1 | GM25953 | Normal | 3 | 0.958 | 0.028 | 0.016 |
| CHR1 | GM25975 | Normal | 3 | 1.009 | 0.036 | 0.021 |
| CHR1 | GM25974 | Normal | 3 | 1.026 | 0.011 | 0.006 |
| CHR1 | GM25952 | Normal | 3 | 0.962 | 0.058 | 0.033 |
| CHR1 | Normal | Normal | 3 | 1.000 | 0.032 | 0.019 |
| CHR4 | GM25953 | Normal | 3 | 1.000 | 0.006 | 0.003 |
| CHR4 | GM25975 | Normal | 3 | 1.000 | 0.012 | 0.007 |
| CHR4 | GM25974 | Normal | 3 | 1.000 | 0.016 | 0.009 |
| CHR4 | GM25952 | Normal | 3 | 1.000 | 0.024 | 0.014 |
| CHR4 | Normal | Normal | 3 | 1.000 | 0.017 | 0.010 |
| CHR8 | GM25953 | Normal | 3 | 1.026 | 0.035 | 0.020 |
| CHR8 | GM25975 | Normal | 3 | 1.027 | 0.053 | 0.031 |
| CHR8 | GM25974 | Normal | 3 | 1.102 | 0.006 | 0.003 |
| CHR8 | GM25952 | Normal | 3 | 1.007 | 0.028 | 0.016 |
| CHR8 | Normal | Normal | 3 | 1.000 | 0.009 | 0.005 |
| CHR10 | GM25953 | Normal | 3 | 0.913 | 0.040 | 0.023 |
| CHR10 | GM25975 | Normal | 3 | 0.998 | 0.024 | 0.014 |
| CHR10 | GM25974 | Normal | 3 | 0.976 | 0.044 | 0.026 |
| CHR10 | GM25952 | Normal | 3 | 0.979 | 0.061 | 0.035 |
| CHR10 | Normal | Normal | 3 | 1.000 | 0.008 | 0.005 |
| CHR12 | GM25953 | Normal | 3 | 0.935 | 0.038 | 0.022 |
| CHR12 | GM25975 | Normal | 3 | 1.094 | 0.005 | 0.003 |
| CHR12 | GM25974 | Normal | 3 | 1.140 | 0.023 | 0.013 |
| CHR12 | GM25952 | Normal | 3 | 1.080 | 0.053 | 0.031 |
| CHR12 | Normal | Normal | 3 | 1.000 | 0.012 | 0.007 |
| CHR17 | GM25953 | Normal | 3 | 0.921 | 0.054 | 0.031 |
| CHR17 | GM25975 | Normal | 3 | 1.061 | 0.061 | 0.035 |
| CHR17 | GM25974 | Normal | 3 | 1.220 | 0.041 | 0.024 |
| CHR17 | GM25952 | Normal | 3 | 1.088 | 0.202 | 0.116 |
| CHR17 | Normal | Normal | 3 | 1.001 | 0.049 | 0.028 |
| CHR18 | GM25953 | Normal | 3 | 0.938 | 0.021 | 0.012 |
| CHR18 | GM25975 | Normal | 3 | 0.991 | 0.028 | 0.016 |
| CHR18 | GM25974 | Normal | 3 | 0.972 | 0.032 | 0.019 |
| CHR18 | GM25952 | Normal | 3 | 0.988 | 0.010 | 0.006 |
| CHR18 | Normal | Normal | 3 | 1.000 | 0.015 | 0.009 |
| CHR20 | GM25953 | Normal | 3 | 0.992 | 0.045 | 0.026 |
| CHR20 | GM25975 | Normal | 3 | 1.104 | 0.007 | 0.004 |
| CHR20 | GM25974 | Normal | 3 | 0.927 | 0.025 | 0.014 |
| CHR20 | GM25952 | Normal | 3 | 0.874 | 0.021 | 0.012 |
| CHR20 | Normal | Normal | 3 | 1.000 | 0.037 | 0.021 |
| CHRX | GM25953 | Normal | 3 | 0.963 | 0.030 | 0.018 |
| CHRX | GM25975 | Normal | 3 | 0.931 | 0.019 | 0.011 |
| CHRX | GM25974 | Normal | 3 | 0.975 | 0.007 | 0.004 |
| CHRX | GM25952 | Normal | 3 | 0.985 | 0.069 | 0.040 |
| CHRX | Normal | Normal | 3 | 1.000 | 0.027 | 0.016 |

**Table S7:** Statistical results for the absolute quantification found in file ANOVA_results.csv. Target Name indicates the genes compared, DF: degrees of freedom, F is the statistic to determine the p-value, MS: mean squares, SS: sums of squares, measure indicates if the tests were dependent measures for example, in a time course, where cell lines were matched across samples. Dist indicates the distribution is normal (parametric).

| Target Name | DF | F | MS | SS | p-value | p-value corrected | Measure | Dist |
| --- | --- | --- | --- | --- | --- | --- | --- | --- |
| GAPDH | 3 | 5.491 | 0.046 | 0.137 | 0.00951 | 0.04753 | dependent | parametric |
| ACTB | 3 | 6.958 | 0.038 | 0.115 | 0.00372 | 0.01859 | dependent | parametric |
| KCNJ6 | 3 | 22.923 | 20.729 | 62.188 | 0.00001 | 0.00004 | dependent | parametric |
| SYP | 3 | 114.917 | 58.478 | 175.433 | 0.00000 | 0.00000 | dependent | parametric |
| GRIA1 | 3 | 11.24 | 10.081 | 30.243 | 0.0004 | 0.00201 | dependent | parametric |

**Table S8:** Post-hoc results from the statistical analysis of the absolute quantification from the one-way ANOVA. These results are found in file Posthoc_result.csv. The comparisons between individual stages for each gene is show. Target Name indicates the gene of interest. **A and B** show the two groups being compared. DF: degrees of freedom, p-value correct is the value corrected for multiple comparisons, p-value before correction for a paired t-test. Parametric, True means a normal distribution was selected.

| Target Name | A | B | DF | p-value corrected | p-value | Paired | Parametric |
| --- | --- | --- | --- | --- | --- | --- | --- |
| KCNJ6 | IPSC | NPC | 5 | 0.85667 | 0.73431 | TRUE | TRUE |
| KCNJ6 | IPSC | DA4W | 5 | 0.00845 | 0.00282 | TRUE | TRUE |
| KCNJ6 | IPSC | DA6W | 5 | 0.00845 | 0.00253 | TRUE | TRUE |
| KCNJ6 | NPC | DA4W | 5 | 0.01157 | 0.00705 | TRUE | TRUE |
| KCNJ6 | NPC | DA6W | 5 | 0.01157 | 0.00771 | TRUE | TRUE |
| KCNJ6 | DA4W | DA6W | 5 | 0.85667 | 0.85667 | TRUE | TRUE |
| SYP | IPSC | NPC | 5 | 0.18543 | 0.171 | TRUE | TRUE |
| SYP | IPSC | DA4W | 5 | 0.0001 | 0.00002 | TRUE | TRUE |
| SYP | IPSC | DA6W | 5 | 0.00018 | 0.00009 | TRUE | TRUE |
| SYP | NPC | DA4W | 5 | 0.00018 | 0.00012 | TRUE | TRUE |
| SYP | NPC | DA6W | 5 | 0.00018 | 0.00011 | TRUE | TRUE |
| SYP | DA4W | DA6W | 5 | 0.18543 | 0.18543 | TRUE | TRUE |
| GRIA1 | IPSC | NPC | 5 | 0.06779 | 0.05649 | TRUE | TRUE |
| GRIA1 | IPSC | DA4W | 5 | 0.03575 | 0.01192 | TRUE | TRUE |
| GRIA1 | IPSC | DA6W | 5 | 0.03575 | 0.01137 | TRUE | TRUE |
| GRIA1 | NPC | DA4W | 5 | 0.06779 | 0.03449 | TRUE | TRUE |
| GRIA1 | NPC | DA6W | 5 | 0.06779 | 0.0519 | TRUE | TRUE |
| GRIA1 | DA4W | DA6W | 5 | 0.35174 | 0.35174 | TRUE | TRUE |

**Table S9:** Example of output from the relative delta-CT analysis from the file clean_data.csv showing the top 10 rows of data. Target Name indicates the gene analyzed, Sample Name indicates the cell line, rq is the relative quantification for each replicate, rq-mean is the mean value of the replicates, rqSD is the standard deviation of the replicates, rqSEM is the standard error of the replicates, Outliers indicates if each outlier is a replicate, Group indicates the group used for statistics for the summary data.

| Target Name | Sample Name | rq | rqMean | rqSD | rqSEM | Outliers | Group |
| --- | --- | --- | --- | --- | --- | --- | --- |
| PAX6 | AIW002-2- | 0.0187 | 0.0223 | 0.0032 | 0.0018 | FALSE | D0 |
| PAX6 | AIW002-2- | 0.0248 | 0.0223 | 0.0032 | 0.0018 | FALSE | D0 |
| PAX6 | AIW002-2- | 0.0235 | 0.0223 | 0.0032 | 0.0018 | FALSE | D0 |
| PAX6 | AIW002-2- | 0.0072 | 0.0073 | 0.0004 | 0.0002 | FALSE | D7 |
| PAX6 | AIW002-2- | 0.0069 | 0.0073 | 0.0004 | 0.0002 | FALSE | D7 |
| PAX6 | AIW002-2- | 0.0077 | 0.0073 | 0.0004 | 0.0002 | FALSE | D7 |
| PAX6 | KYOU-- | 0.1261 | 0.1193 | 0.0065 | 0.0038 | FALSE | D0 |
| PAX6 | KYOU-- | 0.1131 | 0.1193 | 0.0065 | 0.0038 | FALSE | D0 |
| PAX6 | KYOU-- | 0.1187 | 0.1193 | 0.0065 | 0.0038 | FALSE | D0 |
| PAX6 | KYOU-- | 0.0202 | 0.0210 | 0.0007 | 0.0004 | FALSE | D7 |

**Table S10:** Statistical results from the relative quantification comparing the delta-CT and delta-delta-CT using student t-tests. Target name indicate the gene being compared, DF: degrees of freedom, tail; two tail t-test, paired FALSE indicated an unpaired t-test. The p-values are shown under p-val. Model indicates if the delta-CT or the delta-delta-CT method was used.

| Target Name | DF | T | tail | paired | p-value | model | effect size | power | Bayes factor |
| --- | --- | --- | --- | --- | --- | --- | --- | --- | --- |
| PAX6 | 1 | 1.361 | two-sided | FALSE | 0.40342 | delta CT | 1.449 | 0.129 | 0.847 |
| CAMK2A | 1 | -3.277 | two-sided | FALSE | 0.18855 | delta CT | 1.405 | 0.125 | 1.359 |
| GRIN1 | 1 | -3.744 | two-sided | FALSE | 0.16616 | delta CT | 1.836 | 0.162 | 1.454 |
| PAX6 | 1 | 1.361 | two-sided | FALSE | 0.40342 | delta delta CT | 1.449 | 0.129 | 0.847 |
| CAMK2A | 1 | -3.277 | two-sided | FALSE | 0.18855 | delta delta CT | 1.405 | 0.125 | 1.359 |
| GRIN1 | 1 | -3.744 | two-sided | FALSE | 0.16616 | delta delta CT | 1.836 | 0.162 | 1.454 |

**Table S11:** Manual processing compared to Auto-qPCR processing with a range of cut-off values for std to exclude replicates, with or without preserving highly variable outliers. Calculations are all using the absolute model to quantify NRXN3 expression with and without cocaine treatment in three brain regions. Values that differ across processing conditions are highlighted in bold. **Left**, the sample information for Region, Treatment and code name of each mouse (biological replicate) are listed. The processing methods, Manual or Auto-qPCR, are labelled. The std cut-off is the value for which std exceeded for outliers to be moved. The settings for preserving highly variable technical if the ration of mean-media/media is less than 0.1 is indicated by ‘yes’. RNA indicates the RNA quantification values.

|  |  |  | Manual | Auto-qPCR |  |  |  |  |
| --- | --- | --- | --- | --- | --- | --- | --- | --- |
| Preserve high variation replicates |  |  | yes | no | no | no | yes | yes |
| Std cut-off |  |  | 0.29 | 0.3 | 0.275 | 0.2 | 0.3 | 0.275 |
| Region | Treatment | Mouse | RNA | RNA | RNA | RNA | RNA | RNA |
| STN | Saline | B4bis | 0.2564 | 0.2564 | 0.2564 | 0.2817 | 0.2564 | 0.2564 |
| STN | Saline | B6 | 0.1933 | 0.1933 | 0.1933 | 0.1933 | 0.1933 | 0.1933 |
| STN | Saline | R6 | 0.3290 | 0.3290 | 0.3290 | 0.3055 | 0.3290 | 0.3290 |
| STN | Saline | V3 | 0.2845 | 0.2845 | 0.2845 | 0.3357 | 0.2845 | 0.2845 |
| STN | Saline | V4 | 0.3259 | 0.3259 | 0.3259 | 0.3259 | 0.3259 | 0.3259 |
| STN | Cocaine | R5Bis | 0.4570 | 0.4570 | 0.4570 | 0.4570 | 0.4116 | 0.4116 |
| STN | Cocaine | R6bis | 0.1708 | 0.1708 | 0.1708 | 0.1708 | 0.1708 | 0.1708 |
| STN | Cocaine | R8bis | 0.4253 | 0.4253 | 0.4253 | 0.4253 | 0.4253 | 0.4253 |
| STN | Cocaine | V2 | 0.2538 | 0.1659 | 0.1659 | 0.1659 | 0.1987 | 0.1987 |
| STN | Cocaine | V8 | 0.1818 | 0.1818 | 0.1818 | 0.1818 | 0.1818 | 0.1818 |
| GP | Saline | B4bis | 0.2541 | 0.2541 | 0.2541 | 0.2541 | 0.2541 | 0.2541 |
| GP | Saline | R6 | 0.7107 | 0.7107 | 0.7107 | 0.7107 | 0.7107 | 0.7107 |
| GP | Saline | V3 | 0.4125 | 0.4125 | 0.4125 | 0.4125 | 0.4125 | 0.4125 |
| GP | Saline | V4 | 0.2991 | 0.2991 | 0.2991 | 0.2991 | 0.2991 | 0.2991 |
| GP | Cocaine | R5Bis | 0.5021 | 0.5021 | 0.5021 | 0.5021 | 0.4988 | 0.4988 |
| GP | Cocaine | R6bis | 0.9500 | 0.9500 | 0.9500 | 0.9500 | 0.9500 | 0.9500 |
| GP | Cocaine | R8bis | 1.0169 | 1.0169 | 1.0169 | 0.9455 | 1.0169 | 1.0169 |
| GP | Cocaine | V2 | 0.8538 | 0.8538 | 0.8538 | 0.7797 | 0.8538 | 0.8538 |
| GP | Cocaine | V8 | 0.9486 | 0.9486 | 0.9486 | 0.9486 | 0.9486 | 0.9486 |
| SN | Saline | B4bis | 0.7854 | 0.8745 | 0.7854 | 0.7854 | 0.7317 | 0.7317 |
| SN | Saline | R6 | 0.8751 | 1.0784 | 1.0784 | 1.0784 | 0.8751 | 0.8751 |
| SN | Saline | V3 | 1.3306 | 1.3306 | 1.3306 | 1.3306 | 1.1553 | 1.1553 |
| SN | Saline | V4 | 1.0575 | 1.0575 | 1.0575 | 1.0575 | 0.8940 | 0.8940 |
| SN | Cocaine | R5Bis | 1.2379 | 1.2379 | 1.2379 | 1.2379 | 1.2379 | 1.2379 |
| SN | Cocaine | R6bis | 1.0016 | 1.1607 | 1.1607 | 1.2982 | 1.0016 | 1.0016 |
| SN | Cocaine | R8bis | 0.4393 | 0.4393 | 0.4393 | 0.4393 | 0.4393 | 0.4393 |
| SN | Cocaine | V2 | 1.0196 | 1.0196 | 1.0196 | 1.0971 | 1.0196 | 1.0196 |
| SN | Cocaine | V8 | 2.2979 | 2.2979 | 2.2979 | 2.2979 | 2.2979 | 2.2979 |

**Table S12:** Comparison of variance between manual processing and Auto-qPCR. The variance between RNA quantity values calculated manually or with Auto-qPCR were calculated between each mean value found in table S11. For each brain region the sum of the variance was calculated. The same comparison was performed between manual processing and Auto-qPCR with the standard cut-off of 0.3 and the standard cut-off together plus the preserve extreme values option.

| Region | Cut-off 0.3 | Cut-off 0.3 +<br>Preserve |
| --- | --- | --- |
| STN | 0.004 | 0.003 |
| GP | 0.000 | 0.000 |
| SN | 0.037 | 0.030 |
| All regions | 0.037 | 0.033 |

**Table S13:** Addition of group columns used for statistical analysis for one-way and two-way ANVOAs. Treatment (Control, Cocaine), Region (STN,GP,SN) and both together, T_R (STN_Control, STN_Cocaine, GP_Control, GP_Cocaine, SN_Control, SN_Cocaine). Each group column was used for separate one-way ANOVAs. The ‘Treatment’ and ‘Region’ column were used in the two way ANOVA. For visualization the first 15 rows with sample data are shown from the full spreadsheet.

| Well /set | Dye | Content | Description | Efficiency | C(t) | ng | T_R | Treatment | Region |
| --- | --- | --- | --- | --- | --- | --- | --- | --- | --- |
| B1 | FAM | Sample | B4bisNST | 50.57% | 30.71 | 0.128 | STN_Control | Control | STN |
| B1 | TMR | Sample | B4bisNST | N/A | N/A |  | STN_Control | Control | STN |
| B2 | FAM | Sample | B4bisNST | 45.00% | 30.29 | 0.170 | STN_Control | Control | STN |
| B2 | TMR | Sample | B4bisNST | N/A | N/A |  | STN_Control | Control | STN |
| B3 | FAM | Sample | B4bisNST | 46.18% | 30.2 | 0.181 | STN_Control | Control | STN |
| B3 | TMR | Sample | B4bisNST | N/A | N/A |  | STN_Control | Control | STN |
| B4 | FAM | Sample | B4bisGP | 32.11% | 26.6 | 2.053 | GP_Control | Control | GP |
| B4 | TMR | Sample | B4bisGP | N/A | N/A |  | GP_Control | Control | GP |
| B5 | FAM | Sample | B4bisGP | 45.30% | 28.43 | 0.598 | GP_Control | Control | GP |
| B5 | TMR | Sample | B4bisGP | N/A | N/A |  | GP_Control | Control | GP |
| B6 | TMR | Sample | B4bisGP | N/A | N/A |  | GP_Control | Control | GP |
| B6 | FAM | Sample | B4bisGP | 47.44% | 28.18 | 0.708 | GP_Control | Control | GP |
| B7 | TMR | Sample | B4bisSN | N/A | N/A |  | SN_Control | Control | SN |
| B7 | FAM | Sample | B4bisSN | 47.60% | 27.56 | 1.073 | SN_Control | Control | SN |

**Table S14:** Results of the statistical analysis of control vs. cocaine treatment for all brain regions. The results of an unpaired, two tailed students t-test performed in Auto-qPCR using the statistic selections. The table is found in the file ‘ttest_results.csv’.

| Target Name | DF | T | tail | paired | p-val |
| --- | --- | --- | --- | --- | --- |
| B2M | 18.54 | 0 | two-sided | FALSE | 1.000 |
| NRXN3 | 22.74 | -1.555 | two-sided | FALSE | 0.134 |

**Table S15:** Posthoc results after one-way ANOVA comparing brain regions. Control and cocaine treatment samples were pooled together. Target name indicates the gene tested. **A and B** indicate the two regions being compared. DF: degree of freedom.

| Target Name | A | B | DF | p-value corrected | p-value | correction method |
| --- | --- | --- | --- | --- | --- | --- |
| B2M | STN | GP | 9 | 1.0000 | 1.0000 | fdr_bh |
| B2M | STN | SN | 12 | 1.0000 | 1.0000 | fdr_bh |
| B2M | GP | SN | 16 | 1.0000 | 1.0000 | fdr_bh |
| NRXN3 | STN | GP | 9 | 0.0071 | 0.0047 | fdr_bh |
| NRXN3 | STN | SN | 8 | 0.0048 | 0.0016 | fdr_bh |
| NRXN3 | GP | SN | 16 | 0.0549 | 0.0549 | fdr_bh |

**Table S16:** One-way ANOVA and posthoc test comparing groups of brain region and treatment. The ANOVA results are shown for both B2M and NRXN3 for the overall effect of treatment and brain region together (One- way ANOVA. The post-hoc tests for the relevant comparisons are shown for each brain region with and without cocaine treatment (post-hoc).

| Target Name | Comparison | DF | p-value corrected | p-value | Test |
| --- | --- | --- | --- | --- | --- |
| B2M | Treatment and region | 5 | 0.3885 | 0.7771 | One-way ANOVA |
| NRXN3 | Treatment and region | 5 | 0.0006 | 0.0011 | One-way ANOVA |
| NRXN3 | STN_Control vs STN_Cocaine | 8 | 0.9977 | 0.9977 | post-hoc |
| NRXN3 | GP_Control vs GP_Cocaine | 7 | 0.0176 | 0.0334 | post-hoc |
| NRXN3 | SN_Control vs SN_Cocaine | 5 | 0.4127 | 0.4762 | post-hoc |

**Table S17:** Two-way ANOVA and posthoc tests comparing brain region, treatment and interaction. The relevant information was selected from the output files ‘ANOVA_results.csv’ and ‘Posthoc_results.csv’. The 2-way ANOVA results are shown for NRXN3 for the overall effect of brain region (Group1), treatment (Group2) and the interaction effect of region and treatment (Group1*Group2) (upper table). The post-hoc tests for the relevant comparisons are shown for each brain region with and without cocaine treatment for each brain region indicated under contrast. The 2-way ANOVA results are shown on top and the post-hoc multiple t-test comparisons are shown on the bottom, indicated in the Test column

| Target Name | Contrast | DF | p-value corrected | p-value | Test |
| --- | --- | --- | --- | --- | --- |
| NRXN3 | Group1: Region | 2 | 0.0004 | 0.0001 | ANOVA |
| NRXN3 | Group2: Treatment | 1 | 0.2265 | 0.0755 | ANOVA |
| NRXN3 | Group1 * Group2 | 2 | 1.0000 | 0.3513 | ANOVA |
| NRXN3 | all: Control vs Cocaine | 23 | NA | 0.1337 | post-hoc |
| NRXN3 | STN: Control vs Cocaine | 8 | 0.9977 | 0.9977 | post-hoc |
| NRXN3 | GP: Control vs Cocaine | 7 | 0.0529 | 0.0176 | post-hoc |
| NRXN3 | SN: Control vs Cocaine | 5 | 0.6190 | 0.4127 | post-hoc |

